# Diversification dynamics of hypermetamorphic blister beetles (Meloidae): Are homoplastic host shifts and phoresy key factors of a rushing forward strategy to escape extinction?

**DOI:** 10.1101/2021.01.04.425192

**Authors:** E.K. López-Estrada, I. Sanmartín, J.E. Uribe, S. Abalde, M. García-París

## Abstract

Changes in life history traits, including reproductive strategies or host shifts, are often considered triggers of speciation, affecting diversification rates. Subsequently, these shifts can have dramatic effects on the evolutionary history of a lineage. In this study, we examine the consequences of changes in life history traits, in particular host-type and phoresy, within the hypermetamorphic clade of blister beetles (Meloidae). This clade exhibits a complex life cycle involving multiple metamorphoses and parasitoidism. Most tribes within the clade are bee-parasitoids, phoretic or non-phoretic, while two tribes feed on grasshopper eggs. Species richness differs greatly between bee and grasshopper specialist clades, and between phoretic and non-phoretic genera. We generated a mitogenomic phylogeny of the hypermetamorphic clade of Meloidae, including 21 newly generated complete mitogenomes. The phylogeny and estimated lineage divergence times were used to explore the association between diversification rates and changes in host specificity and phoresy, using State-Dependent Speciation and Extinction (SSE) models, while accounting for hidden factors and phylogenetic uncertainty within a Bayesian framework. The ancestor of the hypermetamorphic Meloidae was a non-phoretic bee-parasitoid, and independent transitions towards phoretic bee-parasitoidism or grasshopper specialization occurred multiple times. Bee-parasitoid lineages that are non-phoretic have significantly higher relative extinction rates and lower diversification rates than grasshopper specialists or phoretic bee-parasitoids, while no significant differences were found between the latter two strategies. This suggests that these two life strategies contributed independently to the evolutionary success of Nemognathinae and Meloinae, allowing them to escape from the evolutionary constraints imposed by their hypermetamorphic life-cycle, and that the “bee-by-crawling” strategy may be an evolutionary “dead end”. We show how SSE models can be used not only for testing diversification dependence in relation to the focal character but to identify hidden traits contributing to the diversification dynamics. The ability of blister beetles to explore new evolutionary scenarios including the development of homoplastic life strategies, are extraordinary outcomes along the evolution of a single lineage: the hypermetamorphic Meloidae.

---

Diversification rates (speciation minus extinction) are rarely constant along the evolutionary history of a lineage (Mooers & Heard 1997). Speciation and extinction rates tend to vary over time and among clades as a response to changing abiotic and biotic conditions, such as rapid climate change, geographic range fragmentation, the appearance of a new trait (“key innovation”) or an ecological opportunity resulting from the invasion of a novel niche (Miller 1949; Stanley 1975; Sanderson & Donoghue 1996; Barraclough et al. 1998; Losos & Miles 2002; Ricklefs 2007; Maddison et al. 2007; FitzJohn et al. 2009; Stadler 2011; Rabosky et al. 2013; Donoghue & Edwards 2014; Donoghue & Sanderson 2015; Freyman & Höhna 2019). Shifts in the rate of diversification can have dramatic effects on the evolutionary history of a lineage, and their impact will depend on the magnitude and speed of the change, but also on the interaction with changes in other traits and with the physical template (Vermeij 2001; De Queiroz 2002; Donoghue 2005; Condamine et al. 2018).

The last decade has witnessed the advent of sophisticated methods to infer speciation and extinction rates from phylogenies of extant taxa (Morlon 2014; Sanmartín & Meseguer 2016). Though it is mathematically possible to estimate the extinction rate from a reconstructed time tree including no fossil extinct lineages (Stadler 2011), temporal and clade-specific deviations from constancy in diversification rates leads to inaccurate and often underestimated extinction rates (Rabosky 2010; Morlon 2014). In recent years, different time-dependent and clade-dependent diversification models have been developed to overcome these issues, some assuming continuous time variation while others model change as discrete time steps (Stadler 2011; Morlon et al. 2011; Rabosky et al. 2014; May et al. 2016; Culshaw et al. 2019; Höhna et al. 2019). Nevertheless, concerns that these models lack statistical power and can generate an infinite array of undistinguishable diversification histories remain (Louca & Pennell 2020, but see Morlon et al. 2020).

Recently developed State-Dependent Speciation-Extinction models (SSE) allow researchers to test for a statistical association between the heterogeneity in diversification rates observed within a clade and the rates of evolution of a focal character that is thought to be driving diversification, e.g., a key innovation or ecological opportunity (Maddison et al. 2007; FitzJohn 2010; Beaulieu & O’Meara 2016; Herrera-Alsina et al. 2019; May & Moore 2020). Models such as BiSSE (Binary State Speciation and Extinction, Maddison et al. 2007;) jointly estimate the transition rates among a character states and changes in speciation and extinction rates, and can therefore account for potential interactions between these processes. A perceived advantage of SSE models is that, by explicitly modeling all outcomes in the evolution of a trait with speciation and extinction, they might be able to account for extinct and unobserved lineages and therefore alleviate the problem of undistinguishable diversification histories (Höhna et al. 2019, but see Louca & Pennell 2020). There are, however, warnings about the risk of overconfidence in SEE models, especially the problem of “pseudoreplication” (Maddison & FitzJohn 2015) and inflated Type I error, i.e. an association is detected when there is none. If the phylogeny exhibits high heterogeneity in diversification rates among clades, SSE models may lead to mistaken inference of state-dependent diversification (Rabosky & Goldberg 2015). This issue has spurred the development of Hidden State-dependent Speciation and Extinction models (HiSSE, Beaulieu & O’Meara 2016), allowing for the detected variation in diversification rates to be unrelated to the focal character but instead explained by an unobserved character trait. Typically, the HiSSE model is used to corroborate or reject the association between the rate of evolution of the focal trait and changes in the rate of diversification detected by “standard” SSE models (Condamine et al. 2018; Fernandez et al. 2018; Gajdzik et al. 2019; Nakov et al. 2019). However, these models could also be used to identify the “hidden trait” or unknown causal force whose interaction with the observed trait is behind the heterogeneity in diversification rates; to our knowledge, this other use has never been explored.

A prerequisite for estimating extinction and speciation rates from reconstructed time trees is a sound, well-supported phylogeny. Evolutionary radiations are difficult to tackle with statistical phylogenetic methods because they often exhibit short internal branches, leading to stochastic error, and heterogeneity in gene-tree topologies (due, among others, to incomplete lineage sorting and gene flow), leading to systematic error (Philippe et al. 2011; Cai et al. 2020). In the last decade, the use of genomic approaches has revolutionized phylogenetics, with the possibility to solve radiations in non-model organisms (Barret et al. 2016; Villaverde et al. 2018; Allio et al. 2020; Young & Gillung 2020; but see Cai et al. 2020). In bilaterian animals, including insects, shotgun sequencing of whole mitochondrial genomes has become a favorite tool for reconstructing robust phylogenetic hypotheses at an affordable cost; this is partly due to the small size of the mitochondrial genome, the presence of multiple copies in the cell, and its haploid non-recombinant nature (Finstermeier et al. 2013; Gibb et al. 2016; Yuan et al. 2016; Liu et al. 2019; Yan et al. 2019; Irisarri et al., 2020).

Changes in life history traits, including development of new reproductive strategies and host shifts, are often considered powerful triggers of speciation bursts (Bonett & Chippindale 2004; Hardy & Otto 2014). Host shifts can produce a major turnaround in the evolutionary fate of a parasitic lineage, and might result in a “wild” increase in species number or in new levels of biological complexity (Erwin 1992; Ricklefs & Fallon 2002; Silva et al. 2012). Though host specialization is common in parasites (i.e. the “one-parasite-one host” rule), there is an increasing body of literature showing that changes in host specificity are not infrequent (Hardy & Otto 2014; Nylin et al. 2018; Braga et al. 2020). By jumping hosts, parasites can escape extinction and increase their probability to persist over long evolutionary times (Thines 2019) or over ecological time scales (Brooks et al. 2006; Calatayud et al. 2016). Host shifts have been widely documented among organisms, especially in humans, where the majority of pathogens originate through host changes, including HIV, malaria, and the most recent SARS-CoV-2 (Wolfe et al. 2007; Zhang et al. 2020). Many factors intervene in the evolutionary success of a host shift, including physiological similarity between parasite and host (Runge & Thines 2012; Thines 2019); phylogenetic, ecological and geographical distance between current and potential hosts (Göker et al. 2004; Engelstädter & Fortuna 2019); or differences in parasite and host mutation rates – i.e. higher mutation rates may allow the parasite to overcome host defensive responses – (Gandon & Michalakis 2002). In other words, the likelihood of a new host-parasite interaction and the specific evolutionary outcome of the new relationship are often not determined by a change in a single character (a “key innovation”) but by the interaction of multiple causal agents or changes in different character traits. Changes in different character traits can either appear “simultaneously” during a single speciation event, or as additive factors acting synergistically across several nested speciation events (a “synnovation”, *sensu* Donoghue & Sanderson 2015).

Trait evolution such as host specialization or the development of intricate life-history strategies is deeply affected by the effect of “historical contingency”, defined as “the series of historical details that lead to a specific evolutionary outcome” (Gould 1989). The link between contingency and evolutionary radiation has attracted considerable attention (Losos et al. 1998; Vermeij 2001; De Queiroz 2002; Donoghue 2005; Beatty 2006; Blount et al. 2008; Givnish et al. 2014; Kriebel et al. 2020). According to Beatty (2006), Gould’s contingency” offers two interpretations, which are not mutually exclusive but complementary: a) causal-dependence: a change on a trait is “contingent upon” an earlier change in a different trait, either promoting (De Queiroz 2002; Losos et al. 1998), or restraining it (Vermeij 2001); b) unpredictability: previous states of the character trait are necessary but insufficient to lead to the resulting outcome. So far, contingency in host specialization has never been explored using SSE models. Yet, because host shifts often imply a major change in ecological interactions (Calatayud et al. 2016), they are probably driven by multiple interacting causal factors, which may act in confluence (“synnovation”) or contingent upon one another.

Another interesting consequence of causal-dependence in the context of host-parasite associations is the possibility of becoming an “evolutionary dead-end” (Vamosi et al. 2003). This process is generally associated to the acquisition of a character state with high extinction or low speciation rates, or with irreversibility in transition rates (Goldberg & Igic 2008; Goldberg et al. 2017). Host specialization has sometimes been considered an evolutionary dead-end because the acquisition of a narrow set of food resources (hosts), and concomitant trait adaptations, may limit future diversification, in contrast with phenotypic plasticity in generalist species (Hardy & Otto 2014). Host-jumps that require changes in multiple levels of biological complexity (e.g., morphological, anatomical, physiological, and ecological) are especially difficult to reverse.

Commonly known as “blister beetles”, Meloidae is a family of Coleoptera (Tenebrionoidea), which includes *circa* 3000 described species (Bologna et al. 2008). Its vernacular name is related with the capacity to synthetize cantharidin, a sesquiterpenoid toxin (Percino-Daniel et al. 2013; Bravo et al 2017). This substance is a powerful systemic poison, acting mainly in tissue degradation (lethal dose in humans: 0.5 mg/kg), and a high deterrent for invertebrate and vertebrate predation (Kaiser & Michl 1958; Carrel & Eisner 1974; Bertaux et al. 1988). Cantharidin has also been associated to parasitic regulation in birds (Bravo et al. 2014). Because of the pharmacological usage of cantharidin, blister beetles have drawn scientific attention since the origins of Zoology (Dioscorides 1636; Fischer 1827; Amor Mayor 1860). Meloidae includes three lineages: Eleticinae, with approximately 100 species and a Gondwana-like distribution (absent from Australia); Nemognathinae, comprising ∼520 species and distributed in all continents except New Zealand and Antarctica; and Meloinae, the richest subfamily with almost 2500 species, sharing its geographic distribution with Nemognathinae (Pinto & Bologna 1999; Bologna 2009; Bologna & Pinto 2002). In the most recent phylogeny of Meloidae (Bologna et al. 2008), Eleticinae was reconstructed as sister to the clade formed by Meloinae and Nemognathinae. Eleticinae exhibits the non-parasitic life cycle typical of most Tenebrionoidea, which is characterized by a single event of metamorphosis (Pinto et al. 1996; Bologna & Di Giulio 2011). In contrast, Meloinae and Nemognathinae exhibit a unique, intricate “hypermetamorphic” life cycle. This type of unusual development usually involves three metamorphoses before the imago, with at least four larval phases, each of them so different from each other in morphology and behavior that they could be easily assigned to other families different from Meloidae (Fig. 1; Bologna & Pinto 2001; Bologna et al. 2008).

**Figure 1.**
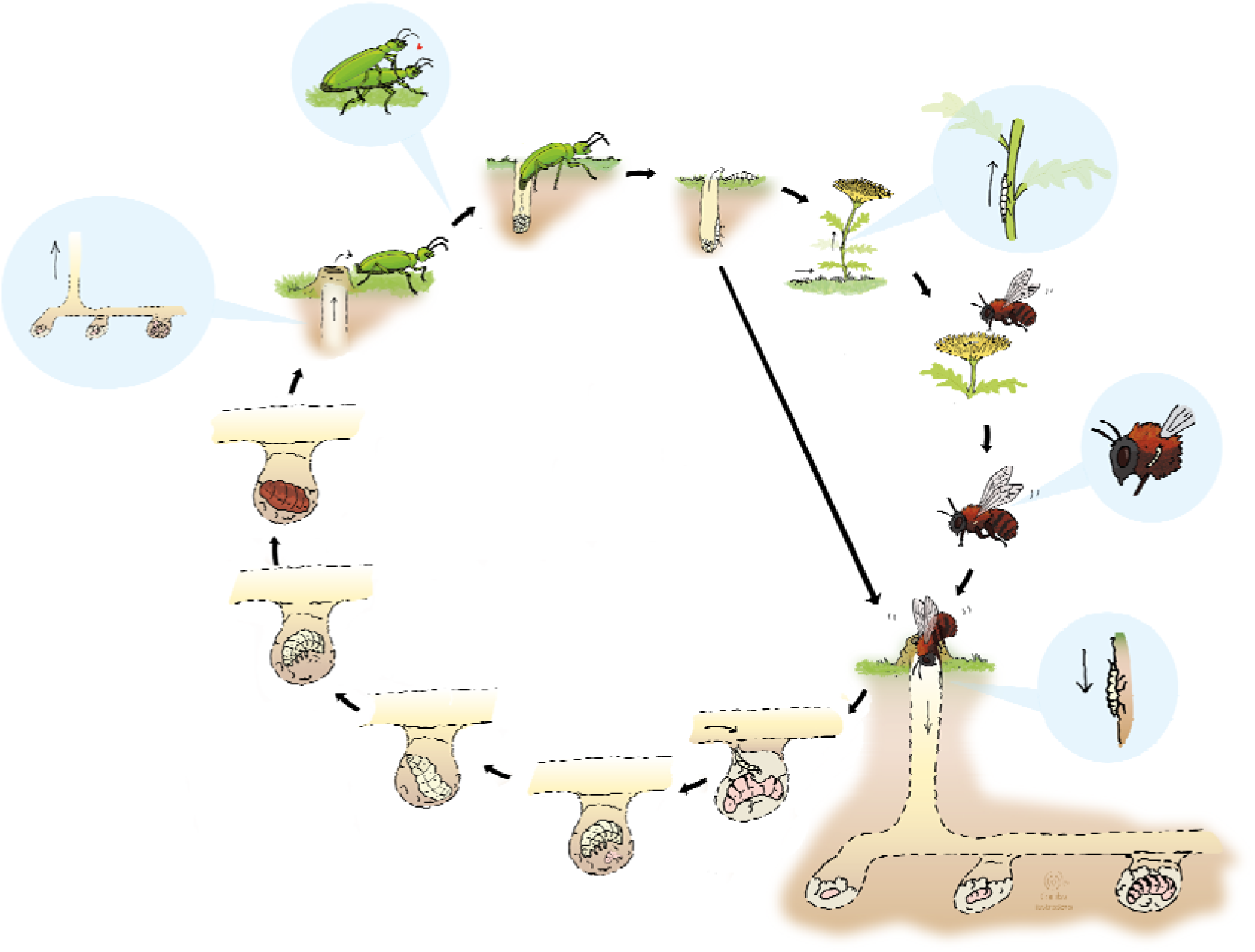
Life cycle of hypermetamorphic Meloidae parasitizing sub-social bees. Eggs are laid in the ground (Meloinae) or on the phyllaries of flowers (Nemognathinae) mainly Asteraceae (Enns 1956). When the eggs hatch, a highly mobile larva emerges from the hatching site and searches for bee nests. This first larva (“triungulin”) can either, depending on the lineage, climb a flower and wait for a bee to visit the flower and then attach to it and be transported to the nest (phoresy), or wander around the ground until it finds an entrance to the bees’ nest (active searching). Once the first larva reaches the bees’ nest, it starts eating the provisions, eggs or larvae of usually a single cell. A first metamorphosis then occurs, and the second larva known as “first grub larva” emerges; it presents reduced motility, but feeds nearly continuously until its metamorphosis. In the second metamorphosis, the first grub larva changes into a “coarctate larva”; the larva loses its appendages and enters into diapause. A third metamorphosis occurs, and a second grub larva morphologically similar to the second larva develops; this larva recovers the motility, although it does not feed. Finally, the larva pupates, and in a few days the adult emerges and the cycle begins again.

Though the hypermetamorphic life cycle is present in all species of Nemognathinae and Meloinae, two traits exhibit variation at the tribal and generic levels: the mode of locomotion used by the first instar larva to reach the food source (Fig. 1), and the host itself (Fig. 2). The four tribes within Nemognathinae and six of the eight tribes included in Meloinae are parasitoids of many different species of solitary or subsocial bees (superfamily Apoidea) feeding on all resources available at the nest: eggs, bee larvae, and provisions (Fig. 1). The two exceptions are Epicautini and Mylabrini, which first instar larvae feed on eggs within the egg-pods of grasshoppers of the family Acrididae (Fig. 2; Bologna 1991; but see Bologna & Di Giulio 2011, for a possible case of parasitizing Sphecidae). Bologna et al. (2008) proposed that Epicautini and Mylabrini were not sister-taxa and as a consequence each event of host-jump to Acrididae was independent from each other (homoplastic). The mode of locomotion of first instar larvae to reach the food source is another trait that varies across taxa; this includes phoresy, i.e. passively latching onto a bee to reach the nest, and active crawling. In phoretic taxa, first instar larvae climb to flowers and attach to passing bees (Fig. 1; Hafernik & Saul-Gershenz 2000); in non-phoretic species, larvae wander on the ground, actively searching for bee nests (Fig. 1) or grasshoppers’ egg-pods (Fig. 2). All species of Nemonagthinae are phoretic parasitoids of bees (with the probable exception of *Stenodera*; Bologna et al. 2002; Bologna & Di Giulio 2011), whereas some tribes of Meloinae that are bee-parasitoids (e.g. Meloini) include phoretic (*Meloe*) and non-phoretic (*Physomeloe*) genera. Phoresy is hypothesized to have evolved at least two times independently in bee-parasitoid Meloinae (Bologna & Pinto 2001; Bologna et al. 2008).

**Figure 2.**
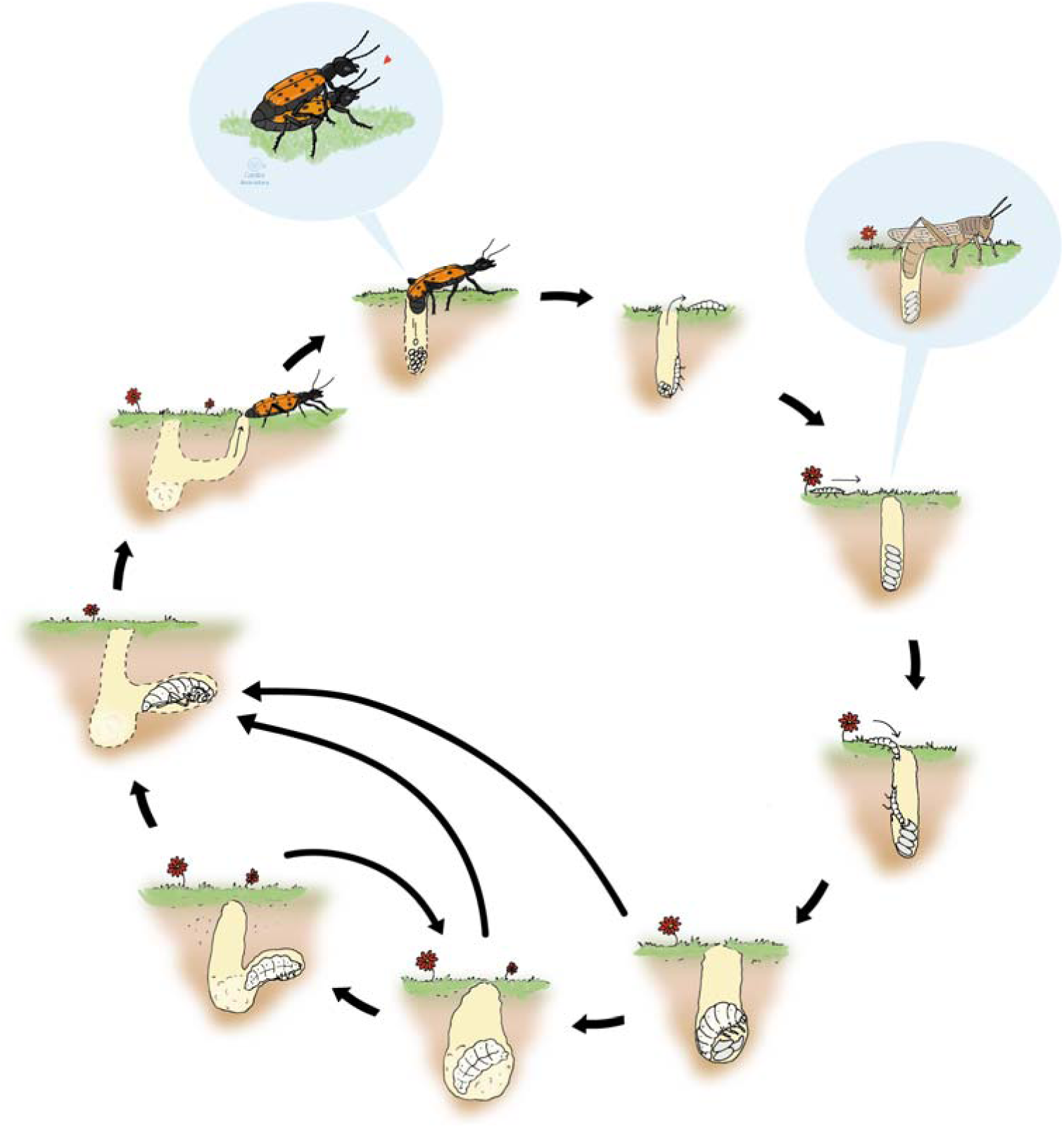
Life cycle of hypermetamorphic Meloidae specialized in grasshopper eggs. Meloid eggs are laid in the ground. A highly mobile larva emerges from the hatching site and actively searches for grasshoppers’ egg-pods. Once the first larva reaches the pod, it starts eating the eggs. A first metamorphosis then occurs, and the second larva known as “first grub fase” emerges. The following larval phases are as those of the lineages that parasitize bees’ nests. Jumps and reversions between larval stages have been observed mainly in Epicautini and Mylabrini (Selander & Mathieu 1964; Selander & Weddle 1969).

Bologna et al. (2008) argued that host specificity could explain the remarkable difference in species richness observed between Nemognathinae (∼500 species), and Meloinae, with ∼2500 species. It may also explain differences among tribes of Meloinae: the two tribes feeding on grasshopper eggs, Mylabrini and Epicautini, include *circa* 600-700 species each, while the largest tribes that parasitize bees (Meloini and Pyrotini) do not exceed 300 species together (Table 1). There is no obvious pattern of species richness across phoretic and non-phoretic genera (Table 1). For example, in the tribe Lyttini, the non-phoretic genus *Lytta* comprises 109 species, while *Lagorina* only includes two. Some phoretic genera of Meloini like *Meloe* are species-rich (153 species), while others (*Spastonyx, Lyttomeloe …*) comprise but a few species (Table 1). Bologna & Pinto (2001) suggested that phoresy could be evolutionarily advantageous in Meloinae, because it allows long-distance dispersal of the first instar larvae, thus potentially increasing the geographic ranges of species; and, second, because transporting the larvae to their food source ensures successful larval rearing. However, these authors also questioned whether this mode of locomotion is more effective in finding the host than the non-phoretic, active crawling behavior present in other genera (Bologna & Pinto 2001). For example, the grasshopper specialist tribes, Mylabrini and Epicautini, are both species-rich and non-phoretic.

**Table 1.** Taxa included in this study, with associated taxonomic diversity, presence of a phoretic behavior, and host-type specialization. Species richness were obtained from Pinto & Bologna (2009), Bologna & Pinto (2002) and Campos-Soldini et al. (2018).

| Family | Tribe | Tribe' species richness | Proportion of Sampled genera | Sampled genus | Genus' species richness | Phoresy | Host |
| --- | --- | --- | --- | --- | --- | --- | --- |
| Meloinae | Epicautini | 564 | 1/5 | *Epicauta* | 492 | no | grasshopper egg pods |
|  | Meloini | 160 | 3/6 | *Meloe* | 150 | yes | bee nest |
|  |  |  |  | *Physomeloe* | 1 | no | bee nest |
|  |  |  |  | *Spastonyx* | 2 | yes | bee nest |
|  | Lyttini | 395 | 4/27 | *Berberomeloe* | 10 | no | bee nest |
|  |  |  |  | *Lagorina* | 2 | no | bee nest |
|  |  |  |  | *Lytta* | 109 | no | bee nest |
|  |  |  |  | *Oenas* | 12 | no | bee nest |
|  | Eupomhini | 26 | 2/8 | *Megetra* | 3 | no | bee nest |
|  |  |  |  | *Tegrodera* | 3 | no | bee nest |
|  | Mylabrini | 735 | 4/11 | *Actenodia* | 20 | no | grasshopper egg pods |
|  |  |  |  | *Croscherichia* | 18 | no | grasshopper egg pods |
|  |  |  |  | *Hycleus* | 450 | no | grasshopper egg pods |
|  |  |  |  | *Mylabris* | 169 | no | grasshopper egg pods |
|  | Pyrotini | 100 | 1/10 | *Pyrota* | 42 | no | bee nest |
| Nemognathinae | Horiini | 15 | 1/3 | *Cissites* | 2 | yes | bee nest |
|  | Nemognathini | 500 | 4/24 | *Apalus* | 21 | yes | bee nest |
|  |  |  |  | *Leptopalpus* | 2 | yes | bee nest |
|  |  |  |  | *Zonitis* | 163 | yes | bee nest |
|  |  |  |  | *Gnathium* | 14 | yes | bee nest |

In this study, we reconstruct phylogenetic relationships and estimate lineage divergence times in the subfamilies Nemognathinae and Meloinae (with special reference to the latter), and use the resulting time tree and macroevolutionary statistical likelihood methods within a Bayesian framework to examine the role played by host-type and phoresy as drivers of diversification in the hypermetamorphic clade of Meloidae.

To date, hypotheses of relationships among major clades of Meloinae and Nemognathinae have been proposed based on morphological characters (Denier 1935; MacSwain 1956; Selander 1964, Kaszab 1969; Bologna & Pinto 2001), or with combined morphological and molecular (*16S* and *ITS2*) datasets (Bologna et al. 2008). These phylogenies, however, exhibit low statistical clade support, and were constructed using traits directly related to the problems we are interested in, i.e. morphological traits associated to phoresy.

To provide an independent robust phylogenetic hypothesis, we sequenced whole mitochondrial genomes for 15 genera of Meloinae, covering 75% representation of the tribal diversity and 20% generic diversity (Table 1). Additionally, we sequenced five genera of Nemognathinae, representing 50% of tribal diversity and 10% generic diversity (Table 1). In total, we generated 21 new mitogenomes by shotgun sequencing. To account for incomplete taxon sampling we developed an integrative approach incorporating the mitogenomic phylogeny, clade species richness, divergence times, and birth-death simulations, to mimic a larger, clade-representative, taxon sampling. We then used state-dependent speciation and extinction (SSE) models that tie changes in diversification rates to transitions between states in a focal character, while accounting for potential interactions with unobserved (hidden) traits (Maddison et al. 2007; FitzJohn 2012; Beaulieu & O’Meara 2016) in a Bayesian framework. Specifically, with this approach we wanted to answer the following questions: (i) are shifts in host-type responsible for the observed difference in species richness among tribes and between subfamilies? (ii) What is the role of phoresy in diversification dynamics? We demonstrate that the heterogeneity in diversification rates observed in the hypermetamorphic clade of Meloidae was the product of a life strategy change, involving either a move towards phoresy or a change of host-type. We further discuss how contingency may have shaped these life strategy changes and how changes in the diversification processes offered new pathways to escape from possible evolutionary dead-ends. Additionally, we show how SSE models can be used to identify an unobserved character, whose interaction with the focal trait is shaping diversification dynamics.

## MATERIALS AND METHODS

### Taxon Sampling, DNA Extractions, Sequencing and Assembly

Our ingroup dataset was composed of 29 mitogenomes of Meloidae (Tenebrionoidea). It included 23 species from 15 genera of Meloinae, representing six out of the eight tribes of the subfamily (Epicautini, Eupomphini, Lyttini, Meloini, Mylabrini, and Pyrotini; representatives of Cerocomini and Tetraonycini could not be included, Table 1), and six species from five genera of the sister subfamily Nemognathinae, representing two of the four recognized tribes (Nemognathini and Horiini; representatives of Palaestrini and Stenoderini were missing; Table 1) (Fig. 3). From this dataset, mitogenomes were newly generated for 21 specimens (Table S1). These specimens were collected in the field, stored in 100% ethanol and deposited at the Museo Nacional de Ciencias Naturales (MNCN-CSIC, Madrid, Spain). Mitogenomes of the remaining eight species were obtained from GenBank. To root the phylogeny, we selected from GenBank ten additional mitogenomes from six families. According to the most recent phylogenetic hypotheses in Coleoptera (Timmermans et al. 2015; Yuan et al. 2016), these ten taxa represent a wide spectrum of the superfamily Tenebrionoidea. A complete list of the specimens included in this study and GenBank accession numbers is shown in Table S1.

**Figure 3.**
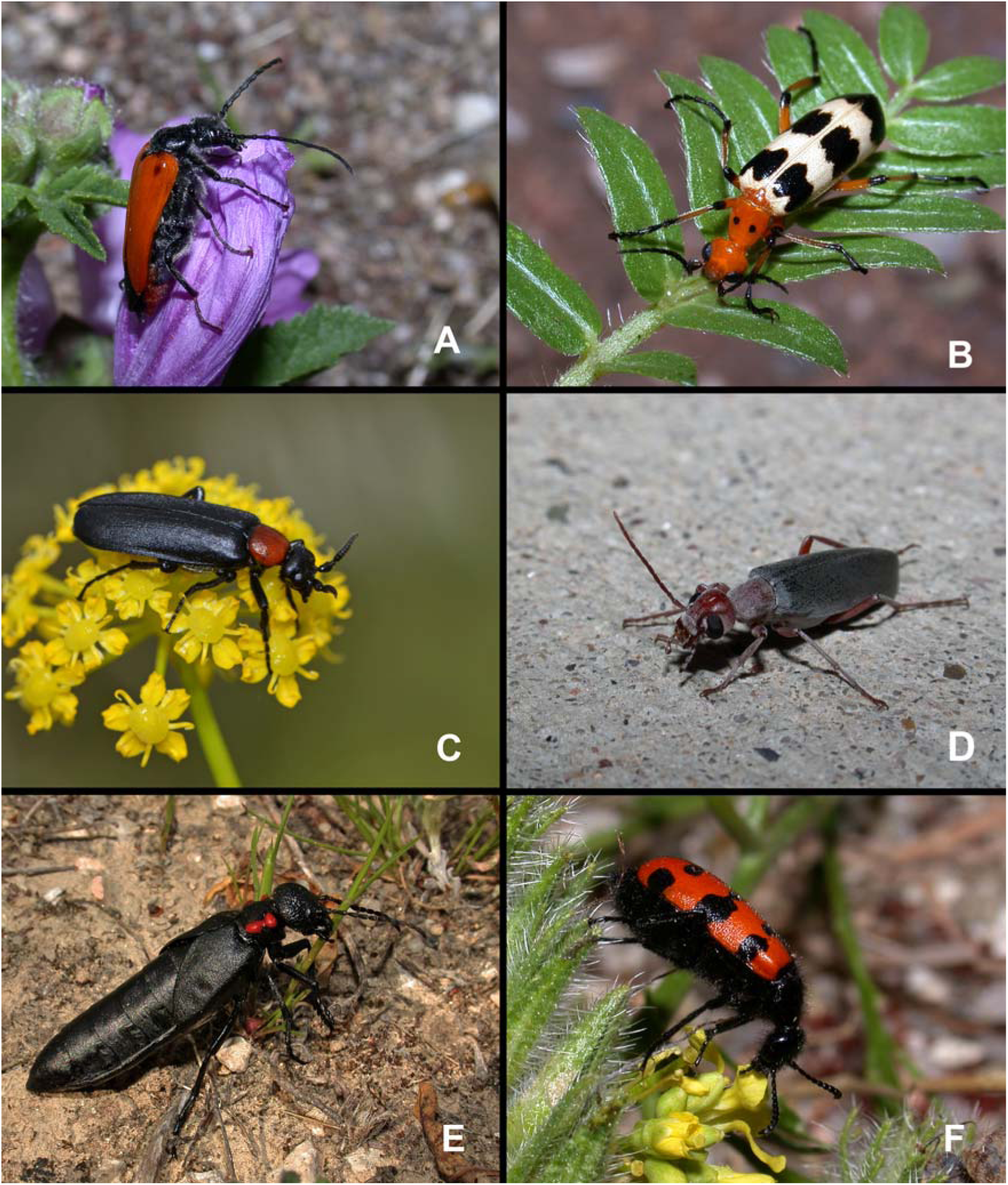
Habitus of representative species of Meloidae used for this study. Live adult specimens of: A) *Apalus guerini* (Mulsant 1858), a phoretic bee specialist (from Perales de Tajuña, Spain) (Nemognathinae: Nemognathini); B) *Pyrota palpalis* Champion 1893, a non-phoretic bee specialist (from Lordsburg, New Mexico) (Meloinae: Pyrotini); C) *Oenas fusicornis* Abeille de Perrin 1880, a non-phoretic bee specialist (from Tielmes, Spain) (Meloinae: Lyttini); D) *Epicauta tenella* (LeConte 1858), a grasshopper specialist (from Needles, California) (Meloinae: Epicautini); E) *Physomeloe corallifer* (Germar 1818), a non-phoretic bee specialist (from Serranillos, Spain) (Meloinae: Meloini); F) *Croscherichia paykulli* (Billberg 1813), a grasshopper specialist (from Moulay Bousselham, Morocco) (Meloinae: Mylabrini). Photographs by MGP.

Tissue samples were obtained from the thoracic muscle of the hind coxae. Total genomic DNA was extracted using the “BioSprint 15 DNA Boold Kit” (Quiagen ®) following the protocol described by the manufacturer. A pair of primers (forward and reverse) were designed in the ribosomal gene *16S* [Mel-16SF and Mel-16SR] (Appendix S1). These two primers, that virtually match in all the *16S* fragments studied in Meloidae, were combined with a set of cox1 pairs of primers (forward and reverse) designed for each tribe. The *16S*-*cox1* primers were used to amplify by long polymerase chain reactions (PCR) the mitochondrial genomes in two overlapped amplicons (with ∼13 kb and ∼5 kb long; the amplification strategy and the sequence of each primer is described in Appendix S1). Amplifications were conducted following the protocol of Uribe et al. (2017). For each mitogenome, two fragments were amplified using the following PCR conditions: denaturing step at 94°C for 60 s; 45 cycles of denaturation at 98°C for 10 s, annealing at 50°C for 30 s, and extension at 68°C for 60 s per kb; final extension step at 68°C for 12 min.

PCR products were cleaned following an ethanol precipitation protocol. Sequencing and assembly schemes were carried out following the protocol described by Abalde et al. (2017). Briefly, an indexed library per sample was constructed using the NEXTERA XT DNA library prep kit (Illumina, San Diego, CA, USA) and sequenced in an Illumina MiSeq platform at Sistemas Genómicos (Valencia, Spain). Mitochondrial genomes were assembled mapping the raw reads back to the *16S* sequences previously Sanger-sequenced, using Geneious ® 11.0.5. We set for the first two iterations a minimum overlap of 60% and a minimum match identity of 99% in order to account for potential sequencing errors. These parameters were increased in the following iterations to 75% and 100%, respectively, until completion of the mitochondrial genomes. Number of reads and mean coverage for each genome are presented in Table S1. Annotation of tRNA genes was carried out through the MITOS2 Web Server (Bernt et al. 2013) and tRNAscan-SE Web Server (Chan & Lowe 2019), which infers cloverleaf secondary structures. The protein-coding genes were checked by aligning them against related mitogenomes to confirm the correct annotation of the start and stop codons. Ribosomal genes were identified and annotated by similarity with the homologous genes of already published mitochondrial genomes of blister beetles and assumed to extend to the boundaries of adjacent genes (Boore et al. 2005).

### Phylogenetic Inference and Divergence Time Estimation

Phylogenetic reconstruction was performed using only protein-coding and ribosomal RNA genes extracted from the complete mitogenomes (Abalde et al. 2017). Sequences of protein-coding genes were extracted into single-gene matrices, subsequently aligned based on the corresponding amino acid translations using TranslatorX Web Server (Abascal et al. 2010), with the MAFFT algorithm (Katoh et al. 2005). Amino acid and nucleotide raw alignments were trimmed with Gblocks (Castresana 2000) using the following specifications: excluding many contiguous non-conserved positions and allowing gap positions within the final blocks. Ribosomal genes were aligned and cleaned through MAFFT and Gblocks online services (Katoh et al. 2017; Talavera & Castresana 2007).

We constructed two datasets using the single-gene matrices: (a) NT-matrix, including all nucleotide sequences (12095 bp) and (b) AA+rNT-matrix, including DNA nucleotide sequences for the non-coding ribosomal genes and the translated amino acid sequences for the coding genes (5170 sites).

Maximum Likelihood (ML) phylogenetic analyses were run on the NT-matrix, using the software RaxML v7.3.1 (Stamatakis 2006) and the “GTRGAMMA” option, which implements the GTR model (Tavaré 1986), with four gamma-distributed rate categories (Yang 1994) for all partitions. ML analyses were conducted with default parameters using the rapid hill-climbing algorithm and 1000 bootstrap pseudo-replicates.

Bayesian Inference (BI) analyses were performed using MrBayes v.3.2.6 (Ronquist et al. 2012). PartitionFinder v2 (Lanfear et al. 2016) was used to select the best partition scheme and molecular evolutionary models for the NT and AA+rNT matrices, under the Bayesian Information Criterion (BIC; Schwarz 1978). MrBayes analyses consisted on two simultaneous chains of 100 million generations each, sampling trees every 10000th generation. Convergence and mixing among chains were evaluated by checking the average standard deviation of split frequencies (< 0.01) and the effective sample size (ESS) values for every parameter (> 200). A majority consensus tree for each analysis was reconstructed after discarding the first 25% of the sampled trees as burn-in.

Differences in biochemical profiles across sites have been shown to be an important source of systematic error in deep-time phylogenomics (Philippe et al. 2011). We used the Bayesian software PhyloBayes (Lartillot et al. 2009) to analyze the NT and the AA (only) data matrices under the site-heterogeneous CAT molecular model: this model implements an infinite mixture model, in which sites are allowed to have different stationary frequencies or biochemical profiles (Lartillot & Philippe 2004). We conducted analyses under the CAT-GTR model, which allows rate variation across sites under the GTR model (Tavaré 1986), and the simpler CAT-Poisson model, assuming a single transition rate for all sequence positions. Analyses were run including both variable and constant sites, since excluding the latter has been shown to mislead phylogenomic inference (i.e. Thode et al. 2020). For every analysis, we ran 4 independent chains of one million generations, sampling every 10th state. After 20000 states, we checked every 5000th generation the convergence of chains. We used the *tracecomp* command to evaluate differences in effective samples sizes (EES) estimated by each chain, with a minimum value of 50 and using a burn-in of 10000 tree samples (100000 generations). Differences in bipartition frequencies between topologies were evaluated with the *bpcomp* and the *readpb* commands; results were summarized using a 10% burn-in, and a sub-sampling of every 10th trees from the posterior distribution. A value equal or lower than 0.1 was used as the maximum accepted difference (*maxdiff*) between chains as a measure of successful convergence, following the PhyloBayes manual (http://www.phylo.org/tools/pb_mpiManual1.4.pdf).

We tested the hypothesis that host shift (bees to grasshoppers) occurred independently in Mylabrini and Epicautini (Bologna et al. 2008). We specified two hypotheses of relationships between Mylabrini and Epicautini to be tested against each other; one involving monophyly of the grasshopper-specialists (e.g. Mylabrini and Epicautini are sister taxa, and consequently host shift occurred once in the history of hypermetamorphic Meloidae) (*H_0_*), and the alternative, in which Mylabrini and Epicautini do not for a monophyletic group (*H_1_*). We used Bayes Factor to compare the marginal likelihood of two models representing each one of the hypotheses obtained applying the topological constraint function in MrBayes to the Bayesian topology (Fig. 4). One model was constructed by constraining the topology reconstructing Epicautini and Mylabrini as a monophyletic group (*H_0_*). For the alternative model (*H_1_*) we used the unconstrained topology (*H_1a_*) (Fig. 4), and also a constrained topology in which Mylabrini was sister to all other Meloinae (including Epicautini) (*H_1b_*).

**Figure 4.**
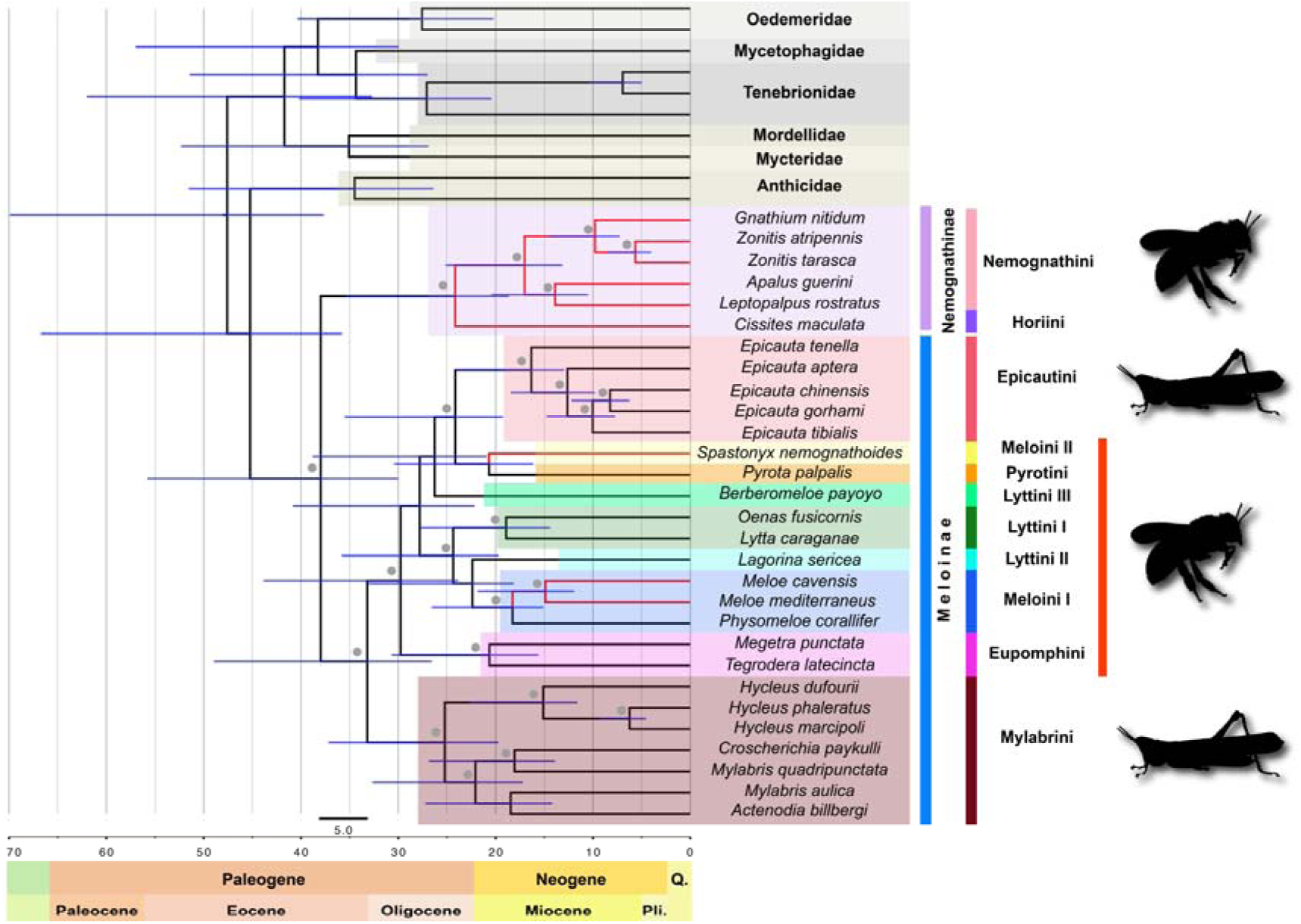
Mitogenomic phylogeny of hypermetamorphic Meloidae. Ultrametric tree obtained using BEAST based on the concatenated mitochondrial dataset. The translated amino acid sequences for the coding genes + the non-coding ribosomal genes matrix (AA+rNT-matrix) was used for this analysis. The chronogram shows mean ages for lineage divergences times using Bayesian relaxed clocks; gray circles near nodes indicate a posterior probability (PP)>0.95 and bootstrap values above 70; purple horizontal bars show 95% HPD values; color shades represent different tribes; red color branches represent phoretic; the characteristic host of each tribe is represented next to its clade.

To estimate the marginal likelihood of each model, we computed power posteriors using the stepping-stone sampling method of Xie et al. (2011) implemented in MrBayes, with the default values for the α–shape parameter of the beta distribution (0.4) and a burn-in of 250000. A MCMC chain of 250000 MCMC steps was sampled every 2500 generations for each of 50 power posteriors, with ß-values ranging between 1 (posterior) and 0 (prior). We compared the two marginal likelihood values using the likelihood ratio test, 2Ln (*H_0_* – *H_1_*), with values larger than 2 indicating positive support for one model over the other, and larger than 6, indicating strong positive support (Kass & Raftery 1995). Since the hypothesis of monophyly (*H_0_*) was rejected, we did not consider necessary adding further constraints to the alternative topology (*H_1a, b_*), because according to Bergsten et al, (2013) the standard Bayes Factor test is biased toward acceptance of the hypothesis of monophyly.

Divergence times were estimated using Bayesian relaxed clocks implemented in BEAST v1.8.2 (Drummond et al. 2012). Analyses were performed using the concatenated AA+rNT dataset. We used for the coding genes the WAG model of amino acid substitution (Whelan & Goldman 2001), while for ribosomal genes we used the HKY model of molecular substitution (Hasegawa et al. 1985). A fossil specimen from the Dominican amber (Cordillera Septentrional, Dominican Republic) was identified and described by Poinar (2009) as a larva of *Meloe dominicanus*. This fossil represents the oldest record for the genus *Meloe* and was used to calibrate the node that clusters the two samples of this genus included in our data set (*M. cavensis* and *M. mediterraneus*). Since the age of Dominican amber is controversial, ranging from 15 to 45 Ma (Cêpek in Schlee 1990; Iturralde-Vincent & MacPhee 1996), we used a log-normal prior distribution with offset=15, mean=10, and standard deviation=1.25 to span the proposed time window for the Dominican amber. Molecular clocks were unlinked across genes, and modeled through an uninformative prior (gamma distribution, initial value=0.01, shape=0.01, offset=0). We used the Birth-Death model with incomplete taxon sampling (Stadler 2009) as the tree growth prior, to account for missing taxa in our dataset. The analysis was run for 100 million generations, sampling every 1000th. We inspected the trace plots and effective sample sizes in Tracer 1.8.0 (Drummond & Rambaut 2007), after discarding the first 20 million generations as burn-in, to assess appropriate convergence and mixing of the chain. All analyses were run in the web public resource CIPRES Science Gateway version 3.3 (Miller et al. 2010).

### State-Dependent Diversification and Incomplete Taxon Sampling

To test for a causal correlation between observed differences in diversity and species variation in traits such as host specificity or mode of locomotion in Meloinae, we used State-dependent Speciation and Extinction (SSE) models (Maddison et al. 2007; FitzJohn 2012; Beaulieu & O’Meara 2016). Incomplete taxon sampling has been shown to severely bias the estimation of diversification rates, especially for the relative extinction parameter (extinction / speciation), under both rate-constant and rate-variable birth-death models (Stadler 2009; Höhna 2014; Morlon 2014; Sanmartín & Meseguer 2016; Louca & Pennell 2020).

Our phylogeny represents less than 1% of the extant species in the family Meloidae. SSE diversification models can potentially account for incomplete taxon sampling by including a parameter *ρ* that incorporates the global sampling fraction, i.e. the proportion of taxa included in the phylogeny relative to the extant diversity (Höhna 2013; Beaulieu & O’Meara 2016). However, this parameter assumes that missing species are uniformly distributed across clades, which is often not the case (Höhna 2014). Ignoring the fact that some clades are better represented than others in the phylogeny, can potentially bias estimates of extinction rates, since the probability that a lineage in state i at time *t* would go completely extinct by the present is 1 minus the sampling fraction, 1– ρ (Maddison et al. 2007; Louca & Pennell 2020). One option to deal with this problem is to distribute incomplete taxon sampling values across clades in the phylogeny, i.e. including clade-specific sampling fractions (Rabosky et al. 2014). However, such a procedure may lead to incorrect estimation of extinction rates in SSE models: artifactual rate shifts in extinction will be inferred wherever any two clades that differ in their sampling fractions coalesce, rendering the likelihood estimation incorrect (Moore et al. 2016; Beaulieu 2020).

To avoid this bias and mimic a larger, clade-representative taxon sampling, which allow us to use SSE diversification models, we developed an integrative approach that incorporates taxonomic information on species richness, the backbone phylogeny, the divergence times previously estimated, and the use of birth-death simulations. First, we calculated the relative extinction rate under a constant-rate diversification model for the original 29-tip phylogeny, using the *bd.shifts.optim* function implemented in the R package *TreePar* (Stadler 2011), with the global sampling fraction set to 0.01%. Second, for several major clades in the backbone phylogeny (tribes, species-rich genera) (Fig. S1), we calculated the net rate of diversification using the *method-of-moments* (MoM) estimator (Magallón & Sanderson 2001), implemented in the function *bd.ms* in the R package *geiger* (Harmon et al. 2008). This function allows estimating the net diversification rate conditioning it to the crown-age of the clade (retrieved from the BEAST MCC tree generated above), the number of extant species (obtained from Pinto & Bologna 1999; Bologna & Pinto 2002; Campos-Soldini et al. 2018), and the relative extinction rate; the latter was fixed to the value estimated by *TreePar* for the backbone phylogeny.

Third, for each of the major clades mentioned above (Fig. S1), we simulated 100 subtrees under a constant-rate model with the function *tess.sim.taxa.age* included in the R package *TESS* (Höhna 2013). Simulations used the MoM estimators of diversification rates for the clade and the *TreePar* turnover value, and were conditioned to clades’ crown age and number of extant species. Fourth, we randomly selected one of the 100 subtrees and bound it to the most inclusive node subtending the clade in the backbone phylogeny, using the *bind.tree* function in the R package *ape* (Paradis et al. 2004). We only simulated subtrees for clades/nodes for which the time of origin was estimated using two species at least. For example, we simulated a subtree with 26 species of Eupomphini (López-Estrada et al. 2019), which was bound to the node connecting the two representative genera *Megetra* and *Tegrodera* in our phylogen*y*; for Nemognathinae, we simulated a 523 species subtree and bound it to the node which was the MRCA of the six species of this subfamily in our backbone phylogeny. Tribes represented by only one tip could not be simulated (e.g. Pyrotini). For some tribes, we simulated subtrees at the generic level, as well, to represent the most speciose genera, for example *Hycleus* in tribe Mylabrini. Figure S1 indicates those nodes in the backbone tree, for which we generated and bound subtrees with associated taxonomic richness. This procedure allowed us to construct a phylogeny with 2150 tips out of approximately 3000 species described for Meloidae, representing 71% of species diversity. It is important to note that these simulated subtrees have no effect on the representation of character states in the phylogeny, as all simulated clades were homogeneous for the trait in question, i.e. each subtree has exactly the same state for all simulated tips. Our aim here was to incorporate the observed differences in species diversity among clades, while decreasing the potential bias introduced by low and uneven sampling when estimating of trait-dependent diversification rates (i.e. subtrees were simulated under clade-specific diversification rates).

To test whether host jump has been a driver of diversification in Meloinae, we used the Binary State-dependent Speciation-Extinction (BiSSE) model (Maddison et al. 2007), allowing for speciation and extinction rates to differ between two states within a character (Maddison et al. 2007; FitzJohn 2010; 2012; Beaulieu & O’Meara 2016; Sanmartín & Meseguer 2016). We used the Bayesian MCMC implementation of this model in the open software RevBayes (Höhna et al. 2016). The hierarchical Bayesian approach to SSE models in RevBayes, represented as directional acyclic graphs (DAGs, Höhna et al. 2014), allows estimation of the marginal posterior probabilities for transition rates and ancestral states through the use of hyperpriors, and is therefore efficient in integrating uncertainty in parameter values (Freyman & Höhna 2019). We coded terminals for two states: parasitoids of bees (0) and grasshopper specialists (1). We followed similar priors to Freyman & Höhna (2019). Speciation and extinction rates for the two states were modeled with a log-normal prior with an expectation centered in the total extant diversity under a constant rate diversification model (rate_mean= ln(Total number of species/2.0)/crown root age); the log-standard deviation was set to a value of = 0.587405 (rate_sd= 0.587405), which assigns a 95% credibility interval that spans one order of magnitude around the mean. Two indirect parameters were also estimated, diversification rate, speciation minus extinction, and relative extinction rate, the ratio of extinction to speciation. Transitions rates were drawn from an exponential distribution with a mean of ten character state transitions over the tree length. Root state probabilities were modeled with a Dirichlet prior with mean= 1. We set the global sampling fraction to 0.71 to account for incomplete taxon sampling in our empirical backbone + simulated subtrees phylogeny. The analysis was run with a chain length of 40000 generations. We summarized ancestral states as nodal marginal posterior probabilities using the maximum a posteriori tree and code provided in RevBayes (https://revbayes.github.io/tutorials/morph/morph_more.html). Finally, we employed a heuristic approximation to stochastic character mapping that does not require a rejection-sampling step (Freyman & Höhna 2019), to estimate the number and timing of transition events between states. Stochastic character mapping was run in RevBayes with 500 time slices. The script to run this analysis is provided in Appendix S2-1.

The frequency of type I error, detecting a signal of trait-dependency when there is not, has prompted some authors to criticize SSE models because of the simplistic, unrealistic null model used for comparison, a lineage evolving with trait-independent but constant diversification rate (Rabosky & Goldberg 2015; Beaulieu & O’Meara 2016). This null model is regarded as a “straw man” because heterogeneity in net diversification rates is to be expected in any phylogeny, except when dealing with very recent times, albeit not necessarily correlated with the focal trait (Rabosky & Goldberg 2015). This issue spurred the development of “hidden-trait” SSE models, such as the Hidden State Speciation Extinction (HiSSE) model (Beaulieu & O’Meara 2016), which allows for the detected variation in diversification rates to be unrelated to the focal character but instead explained by other unobserved character. The hidden, uninformative trait interacts with the character of interest and acts as a “stand-in” for the unknown evolutionary factor that is leading the observed heterogeneity in diversification rates (Beaulieu & O’Meara 2016). Our HiSSE model contained two hidden states (A, B) within each of the two observed states of “host-type”; in total, four character states: parasitoids of bees (0) with hidden states A and B (resulting in 0A and 0B), and grasshopper specialists (1) with hidden states A and B (1A and 1B). If the differences in speciation and extinction rates detected by BiSSE for the observed character states 0 and 1 holds for the two hidden states (A and B), we can conclude that the focal character drives diversification rate heterogeneity among clades. When this is not the case, the observed differences in species richness among clades cannot be associated to the focal character, but to a hidden trait to which diversification is related (Freyman and Höhna 2019).

Initial runs with the same priors as BiSSE resulted in poor mixing. We used instead alternative priors suggested in the RevBayes tutorial (https://revbayes.github.io/tutorials/sse/hisse.html). Speciation and extinction rates for the observed states were modeled as an identical broad log-uniform distribution with bounds “1E-6 and 1E2”; hidden states were modeled through a discretized log-normal distribution, with the number of categories (quantiles) equal to the number of hidden states; the mean of the log-normal for each hidden rate was set to 1.0, to make them relative, and the standard deviation set to an exponential distribution with mean= 0.587405. Transitions rates between observed states were allowed to differ and modeled through an exponential distribution prior with lambda 10, as in the BiSSE model; transition rates between hidden sates were constrained to be equal. All other prior settings followed BiSSE. The analysis was run for 40000 generations, and results summarized as above. Stochastic character mapping was used in RevBayes to obtain marginal probabilities for the number and timing of transitions events between the four-joint observed*hidden states, using 500 time slices. The script to run this analysis is provided in Appendix S2-2.

To test for the potential effect of phoresy and host-type on the diversification rates of hypermetamorphic Meloidae, we ran a second analysis using the Multiple State-dependent Speciation and Extinction (MuSSE) model (FitzJohn 2012). We considered three character states: (0) “bee-by-phoresy”: phoretic parasitoids on bees, i.e. lineages that feed on eggs, larvae and provisions of the bees and use adult bees as a mean of transport to the nest; (1) “bee-by-crawling”: non-phoretic parasitoids of bees, i.e. lineages that feed on eggs, larvae and provisions of the bees but use crawling on the ground to find bee nests; (2) “grasshopper-by-crawling”: non-phoretic lineages that feed on acridid eggs and use crawling to find the egg pod. Since phoresy only occurs in bee-parasitoids, it is not possible to study the joint evolution of phoresy and host-jump as two independent binary traits: i.e. there are no lineages that are “phoretic grasshopper specialists”. Settings for this analysis were similar to the BiSSE analysis, with log-normal priors for speciation and extinction rates and transitions between observed states modeled as an exponential distribution centered on a mean of 10 transitions over the tree length. A pre-burnin step of 5000 generations was used for parameter auto-tuning before a final MCMC chain length of 20000 generations. We also performed stochastic character mapping and estimation of marginal probabilities for ancestral states in the maximum a posteriori tree using the same procedure as BiSSE. The script to run this analysis is provided in Appendix S2-3.

Second, we tested the robustness of the MuSSE results to Type I error or “false positives”, i.e. a trait different from changes in phoresy and host-type is leading rate heterogeneity among lineages. We ran a multiple-state HiSSE analysis (MuHiSSE), with two hidden states (A, B) associated to each of the three observed states in MuSSE (0, 1, 2); in total, the model included six states (0A, 1A, 2A, 0B, 1B, 2B). We used the same priors as in the HiSSE model above, with a log-uniform distribution for observed speciation and extinction rates, and a discretized log-normal distribution for the hidden speciation and extinction rates. A pre-burnin step of 10000 generations was used for auto-tuning before running the final MCMC chain with 40000 generations. Because mixing was difficult, we ran two analyses in parallel and merged the MCMC posterior probabilities of the two runs to increase sample size (i.e. reported results are based on this combined run). The script to run this analysis is provided in Appendix S2-4.

Both standard SSE models (BiSSE, MuSSE) and hidden SSE models (HiSSE, MuHiSSE) assume that the heterogeneity in diversification rates among clades is structured in the phylogeny as an evolving trait, i.e. as discrete shifts. When this is not the case, for example, rates of diversification increasing continuously in one clade and decreasing in another clade, as a response to abiotic factors, HiSSE models can be affected by “false positives”, as well (i.e. accepting trait-dependence when none exists (Rabosky & Goldberg 2017). To provide a different null model to compare against the MuSSE model, we ran a Character-Independent Diversification (CID) model (Caetano et al. 2018). In the CID model, speciation and extinction rates among the states of the focal trait are constrained to be the same, but they are allowed to vary among the hidden states, in other words, rate-heterogeneity is part of the model but is unlinked to the focal observed character (Caetano et al. 2018). Our CID model had two hidden states (A and B) within each of the observed character traits (0, 1 and 2) (CID2). Thus, the model included six transition rates, as well as two speciation and two relative extinction rates, corresponding to the two hidden states (those of the observed focal states are assumed to be equal): 0A=0B, 1A=1B, 2A=2B.

Figures S2-S6 represent each of the models ran in this study (BiSSE, HiSSE, MuSSE, MuHiSSE, and CID2) as DAGs, indicating the parameter dependencies and prior distributions; DAGs were plotted with the graphical software GraphViz (Ellson et al. 2004). We compared the fit of each model to the data using Bayes Factor comparisons of the model marginal likelihood, which were estimated via path sampling and stepping-stone sampling using parallel power posterior analyses in RevBayes (Höhna et al. 2016). We ran 100 power posteriors, with a pre-burnin of 10000 generations, and a 1000-generation chain-length for each power posterior run. The script to run this analysis is provided in Appendix S2-5.

Finally, we tested the robustness of our results against phylogenetic uncertainty, that is, the effect of choosing a particular random simulated subtree to represent clade diversity in the empirical-simulated phylogeny. First, using an R loop script (Appendix S2-6), we generated 100 additional empirical-simulated phylogenies, with alternative random subtrees selected and bounded for clades in the backbone phylogeny. We then checked that all simulated phylogenies belonged to the same “congruence diversification class” (Louca & Pennell 2020) by calculating their “pulled speciation rate” (PSR) with the *fit_hbd_psr_on_grid* function, included in the R package *castor* (Louca & Doebeli 2018); we confirmed that all simulated trees shared similar PSR mean values, allowing for a mean relative deviation of 0.5% (Louca & Pennell 2020). Second, we compared posterior estimates of state-dependent speciation and extinction rates across the simulated phylogenies to ensure differences were not dependent on the topology/branch lengths of the selected random subtrees. We ran a MuSSE analysis, with the same settings as above, over the distribution of 100 simulated phylogenies; we used the multiple-processor *mpi* version of RevBayes (Höhna et al. 2016) and a Unix bash shell script to perform a loop across phylogenies and to record the results. We then performed pairwise comparisons of the posterior distributions of the speciation, extinction, net diversification, and relative extinction across the 100 simulated-empirical trees. If the estimated difference was centered on a mean of 0, we concluded that there was no significant difference in our MuSSE estimates due to the random choice of simulated subtrees, i.e. there is not phylogenetic uncertainty effect.

State-dependent analyses were performed on the Hydra supercomputer provided by the facilities of the Laboratories of Analytical Biology (LAB) of the National Museum of Natural History, Smithsonian Institution. Appendix S1 and S2 are available at https://github.com/isabelsanmartin/Trait-dependent-analyses-Meloinae

## RESULTS

### Sequencing, Assembly and Mitogenome Organization

A total of 14 complete and 7 partial mitogenomes of 21 species of Meloidae were obtained. The number of reads, mean coverage, and length of each mitogenome are provided in Table S1 and Appendix S1. Genome organization was shared across all complete sequenced mitogenomes, and followed the molecular organization described by Du et al. (2016; 2017). The circular genome, represented in Figure S7, encoded for 13 protein-coding genes, and 2 rRNAs and 22 tRNA genes, and also contained a putative control region. Major strand encodes genes: *cox1*, *cox2*, *cox3*, *cytB*, *ATP6*, *ATP8*, *NAD2*, *NAD3*, and *NAD6*; as well as the following tRNAs: *L2*, *K*, *D*, *G*, *A*, *R*, *N*, *S1*, *E*, *T*, *S2*, *I*, *M* and *W*. Minor strand encodes genes: *NAD1*, *NAD4*, *NAD4L*, *NAD1* and ribosomal *16S* and *12S*; as well as the following tRNAs: *F*, *H*, *P*, *L1*, *V*, *Q*, *C* and *Y*.

### Phylogenetic Inference and Divergence time estimation

Phylogenetic inference was obtained using the NT-matrix or the AA+rNT datasets, and different inference methods: Maximum Likelihood in RAxML under site-homogeneous models (Fig. S8), Bayesian Inference in MrBayes under site-homogeneous models (Fig. S9), and Bayesian inference under relaxed clock models in BEAST (Fig. 4). The topology exhibiting higher posterior probability values (PP) was obtained with Bayesian Inference in MrBayes and the AA+rNT dataset (88% of nodes exhibited values equal to 1) (Fig. S9). Bayesian inference with site-heterogeneous models in PhyloBayes did not converge for the more complex CAT-GTR model, due to poor mixing (results not shown). PhyloBayes results under the simpler CAT-Poisson analysis are shown in Fig S10; this tree was largely congruent with the ML, MrBayes and BEAST topologies above, recovering all major clades; however, resolution was considerably lower, especially for intertribal relationships (PP < 0.5).

Subfamilies Nemognathinae and Meloinae were recovered as reciprocally monophyletic (PP =1/ BS =100) (Fig. 4). Within Nemognathinae, the genus *Cissites*, representing the tribe Horiini, is placed as sister to all included representatives of Nemognathini (1/100). The first splitting event within Meloinae separates the tribe Mylabrini, recovered as monophyletic (1/100), as sister to a clade that includes all remaining taxa (1/89). Within this larger clade, relationships among tribes and genera were not fully resolved (0.6/28, Fig. 4). Tribes Eupomphini and Epicautini are also recovered as monophyletic (1/100).

Meloini was found to be non-monophyletic. Species belonging to *Meloe* and *Physomeloe* form a clade (Meloini I) (1/100), closely related to *Lagorina sericea* (Lyttini II), and sequentially, to a clade that includes *Oenas fusicornis* and *Lytta caragenae* (LyttiniI) (1/100). In contrast, *Spastonyx nemognathoides* (Meloini II) was more closely related to Pyrotini and Epicautini than to Meloini I. These two subclades, Meloini I-Lyttini I-Lyttini II and Epicautini-Pyrotini-Meloini II, received high support in the Bayesian analyses, but not in the maximum likelihood analyses (1/43; 1/40, respectively). Lyttini was also recovered as non-monophyletic. Representative species and genera were arranged in three non-sister lineages: Lyttini I and Lyttini II, as mentioned above, and *Berberomeloe payoyo* (Lyttini III), sister to the subclade Epicautini-Pyrotini-Meloini II, albeit with low clade support (Fig. 3).

Bayes Factor comparison between the marginal likelihoods of the unconstrained and constrained analyses rejected the monophyly hypothesis (*H_0_*) for Epicautini plus Mylabrini in favor of the alternative hypothesis (*H_1_*), with 2lnBF= 2*((-93393.74)-(-93407.19))= 26.9, which, according to the scale given in Kass and Raftery (1995), can be interpreted as very strong support against a monophyletic grasshopper specialists’ group (Epicautini plus Mylabrini).

Estimates of lineage divergence times with relaxed molecular clocks in BEAST (MCC tree, Fig. 4) dated the crown-age of the family Meloidae in the Eocene (Mean 37.97 Ma, 95% HPD 29.99–55.81 Ma). An Eocene-Oligocene origin was estimated for the MRCA of the subfamily Meloinae (Mean 33.19 Ma, 26.57–48.94 Ma), whereas the MRCA of Nemognathinae was inferred as Late Oligocene-Early Miocene (Mean 24.16 Ma, 18.64–35.43 Ma). Mylabrini originated in the Late Oligocene (Mean 25.25 Ma, 19.72–37.19 Ma; the MRCAs of the remaining tribes ranged between 20 and 14 Ma) (Fig. S11).

### State Dependent Diversification

State-dependent Speciation and Extinction Diversification analysis with BiSSE (Fig. 5) supported higher speciation and extinction rates in bee-parasitoids (state 0) compared to those shown by clades feeding on grasshopper eggs (1) (Fig. 5). Net diversification rates were also lower, and relative extinction rates higher, for bee parasitoids than for grasshopper specialists. There was some overlap between the marginal posterior distributions of speciation and extinction rates (Fig. 5). However, pairwise comparisons of values across the MCMC posterior set produced a distribution of differences (s0–s1) in which the 95% credibility interval was larger or smaller than zero (i.e. versus overlapping zero), indicating significant differences between speciation rates or between extinction rates (bee vs grasshopper specialists) (Fig. S12). Bayesian reconstruction of ancestral states in the maximum a posteriori tree (Fig. 5) showed that the ancestral condition for the MRCA of Nemognathinae and Meloinae was bee-parasitoid. Two independent events of host-jump from bees towards grasshoppers, in tribes Mylabrini and Epicautini, were recovered by stochastic character mapping analysis (Fig. S13).

**Figure 5.**
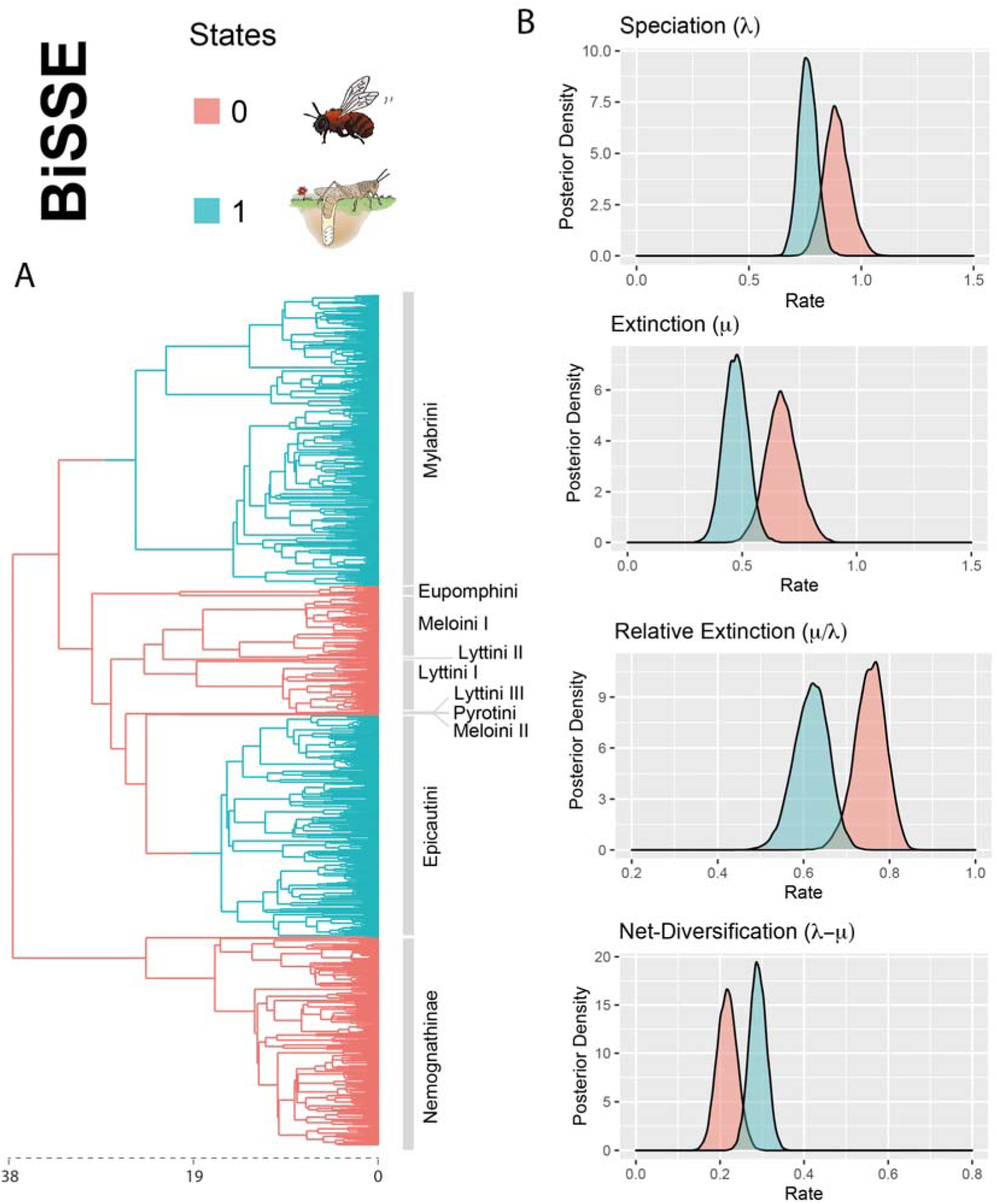
Maximum a posteriori reconstruction of host choice evolution in Meloidae and trait-dependent posterior distributions of diversification rates estimated through BiSSE. (A) Host choice evolution simulated under Bayesian stochastic character mapping; divergence times in millions of years are indicated by the axis at the bottom of the tree; branch colors denote different host; transitions between character states are indicated by changes in color along the branches; note that the state 0 (bee-parasitoid) is reconstructed as the ancestral state of the hypermetamorphic Meloidae and also as the ancestral state of each family. (B) Posterior densities of speciation (λ), extinction (μ), relative extinction (μ/λ) and net-diversification (λ−μ) rates. Colors correspond to the posterior probabilities for a given state; changes in host choice from state 0 to 1 are associated with diversification rate heterogeneity. Bee parasitoids lineages show higher speciation and extinction rates than grasshopper specialists, thus grasshopper specialists’ lineages are associated with the higher diversification rates and lower relative extinction rates than bee parasitoids.

The HiSSE model, allowing for the existence of correlated hidden traits (Fig. 6) supported a somewhat different pattern: differences in rates of speciation were still present between the two observed states, 0 and 1, but not for extinction. As in the BiSSE analysis, the net diversification rate and the rate of relative extinction were lower and higher, respectively, for bee-nests parasitoids than for grasshopper specialists, and these differences were maintained within each hidden state (Fig. 6). However, differences for these rates were larger for the hidden states than for the observed states, with state B showing higher speciation rates than state A (Fig. 6). In fact, for some parameters there was overlap between the marginal posterior distributions of opposite observed*hidden states, such as 1A and 0B, indicating a strong effect of the hidden trait (Fig. 6). Reconstruction of ancestral states (Fig. S14a) and stochastic character mapping of transition events in the maximum a posteriori tree (Fig. 6, Fig. S14b) suggested that the “hidden trait” was phoresy: lineages that behave as phoretic parasitoids of bees were reconstructed as 0B (e.g. Nemognathinae, *Meloe*), while non-phoretic parasitoids of bees were inferred as 0A (e.g. Pyrotini, Eupomphini). The state of the most recent common ancestor (MRCA) of subfamilies Nemognathinae and Meloinae was reconstructed as 0A, as well as the MRCA of each subfamily (Fig. 6). Intriguingly, the ancestor of genus *Cissites*, a phoretic parasitoid on bees, as well as the ancestor of the remaining Nemognathinae, were reconstructed as 0A, with this state changing to 0B in the terminal part of the branch (Fig. 6). However, marginal posterior probabilities of character states were low on the longest branches, such as the one subtending *Cissites*, indicating uncertainty in the reconstruction (Fig. S13b).

**Figure 6.**
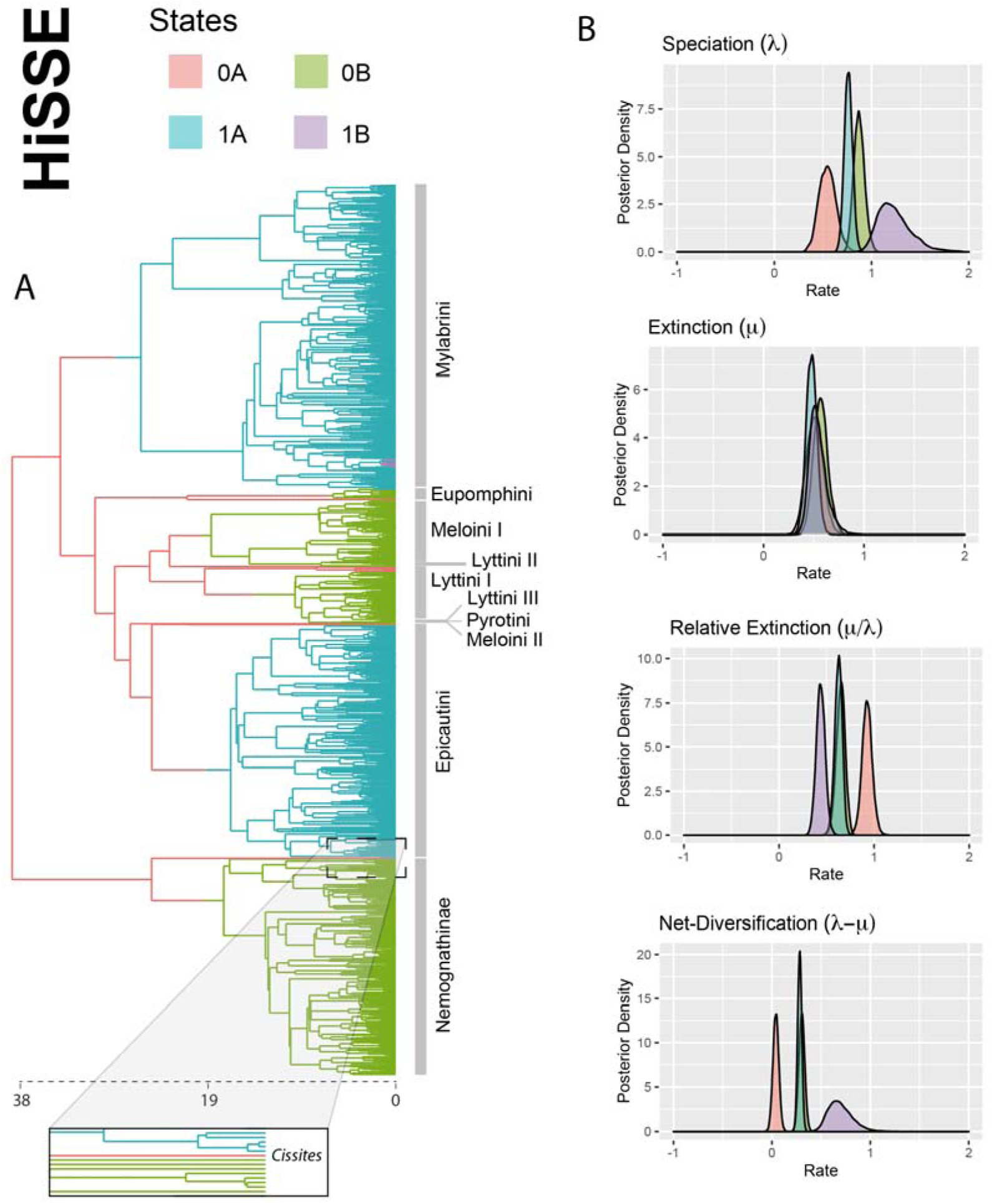
Maximum a posteriori reconstruction of host choice evolution in Meloidae and trait-dependent posterior distributions of diversification rates estimated through HiSSE. (A) Host choice evolution simulated under Bayesian stochastic character mapping; divergence times in millions of years are indicated by the axis at the bottom of the tree; branch colors denote the four different states, being 0 and 1 the observed states (bee parasitoid and grasshopper specialists, respectively) and A and B the hidden states; transitions between character states are indicated by changes in color along the branches; note that lineages reconstructed as 0B coincide in most cases with the phoretic lineages, such as Nemognathinae and Meloini II, while non-phoretic parasitoids of bees, were reconstructed as 0A such as Lyttini II and Pyrotini; inset panel is signaling that the phoretic genus *Cissites* was reconstructed as 0A while the rest of the subfamily Nemognathinae was reconstructed as 0B. (B) Posterior densities of speciation (λ), extinction (μ), relative extinction (μ/λ) and net-diversification (λ−μ) rates. Colors correspond to the posterior probabilities for a given state; note that the posterior densities of speciation, relative extinction and diversification rates are in partial agreement with BiSSE results, however, the overlapping of the marginal posterior distributions for states 1A and 0B indicates that the background rate changes are unassociated with the trait in question: host-type.

The importance of phoresy to explain differences in diversity related to host specificity in Meloidae was confirmed by the MuSSE analysis (Fig. 7). Significant differences in speciation and extinction rates were found between bee-parasitoids and grasshopper specialists, but only for the non-phoretic lineages. Non-phoretic parasitoids of bees (state 1) exhibited marginally higher speciation rates and significantly higher extinction rates than grasshopper specialists (state 2), and also than phoretic parasitoids of bees (state 0), resulting in significantly lower diversification rates and higher relative extinction rates for state 1 than for states 0 and 2 (Fig. 7). In contrast, no significant differences were found between phoretic bee-parasitoids and grasshopper specialists’ lineages, with overlapping marginal posterior distributions for states 0 and 2 across all parameters (Fig. 7). Reconstruction of ancestral states (Fig. S15) and stochastic character mapping (Fig. 7) supported non-phoretic bee-parasitoid as the ancestral state for the MRCA of Nemognathinae and Meloinae. Transition events from non-phoretic bee-parasitoids towards phoresy in bee-parasitoids, and towards a new grasshopper-host, took place in hypermetamorphic Meloidae in at least five independent events (Fig. 7). Introducing hidden traits in this model (MuHiSSE) did not change these results (Fig. 8), with hidden state B showing larger differences among the focal states than hidden state A. Likewise, our inferences were robust to the choice of subtrees for the empirical backbone-simulated phylogeny in the MuSSE analysis; pairwise comparisons across the 100 empirical-simulated phylogenies show a distribution of values with a mean centered on 0 (Fig. S16).

**Figure 7.**
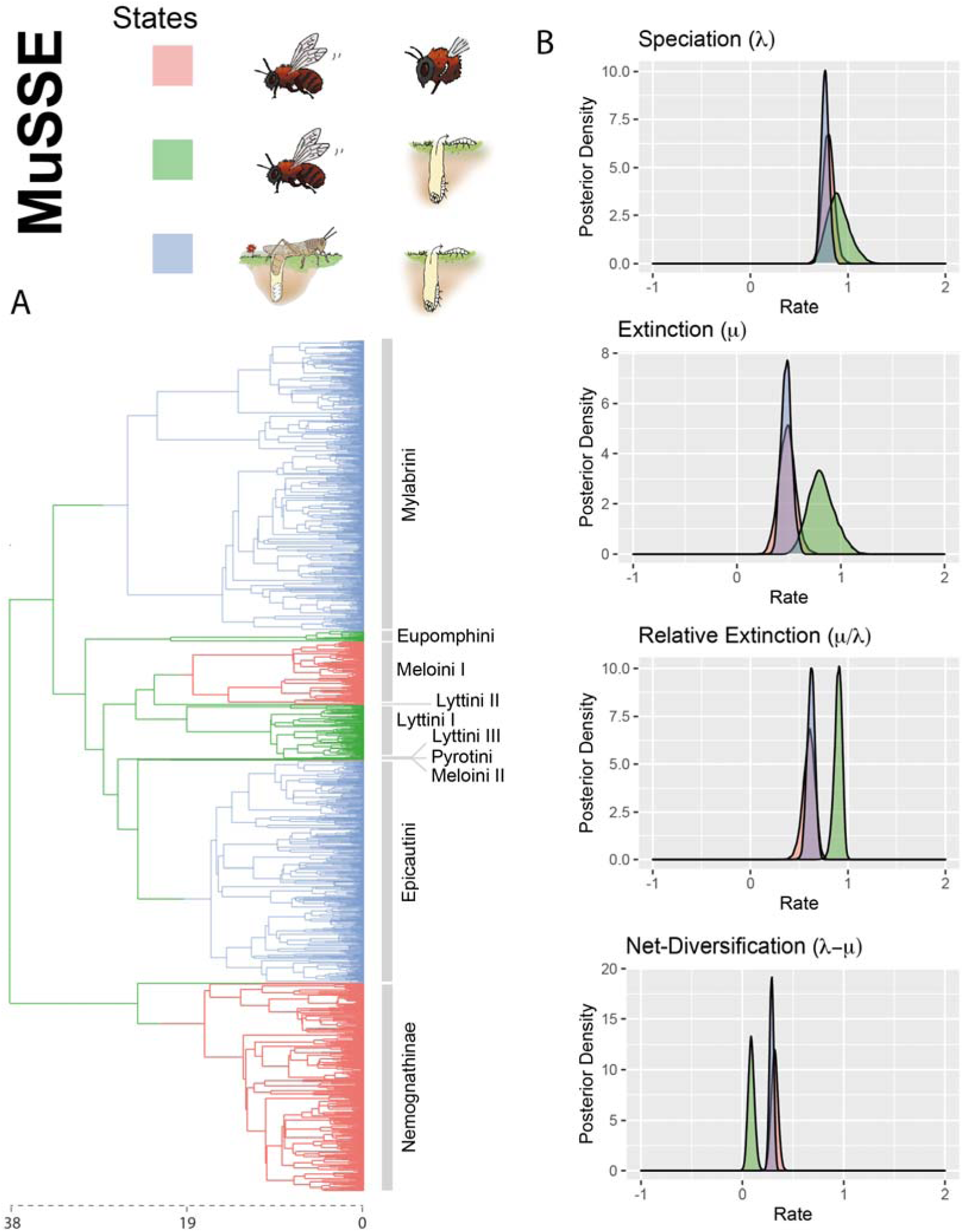
Maximum a posteriori reconstruction of host choice evolution in Meloidae and trait-dependent posterior distributions of diversification rates estimated through MuSSE. (A) Life strategy evolution simulated under Bayesian stochastic character mapping; divergence times in millions of years are indicated by the axis at the bottom of the tree; branch colors denote different life strategies; transitions between character states are indicated by changes in color along the branches; note that a non-phoretic parasitoids of bee nest is reconstructed as the ancestral state of the hypermetamorphic Meloidae and also as the ancestral state of each family. (B) Posterior densities of speciation (λ), extinction (μ), relative extinction (μ/λ) and net-diversification (λ−μ) rates. Colors correspond to the posterior probabilities for a given state; changes in life strategy are associated with diversification rate heterogeneity. The ancestral non-phoretic parasitoids of bees’ nests showed the lowest diversification rates and the highest relative extinction rates, while there was no significant difference neither in diversification rates nor relative extinction rates between phoretic parasitoids of bee nest and parasitoids of grashoppers.

**Figure 8.**
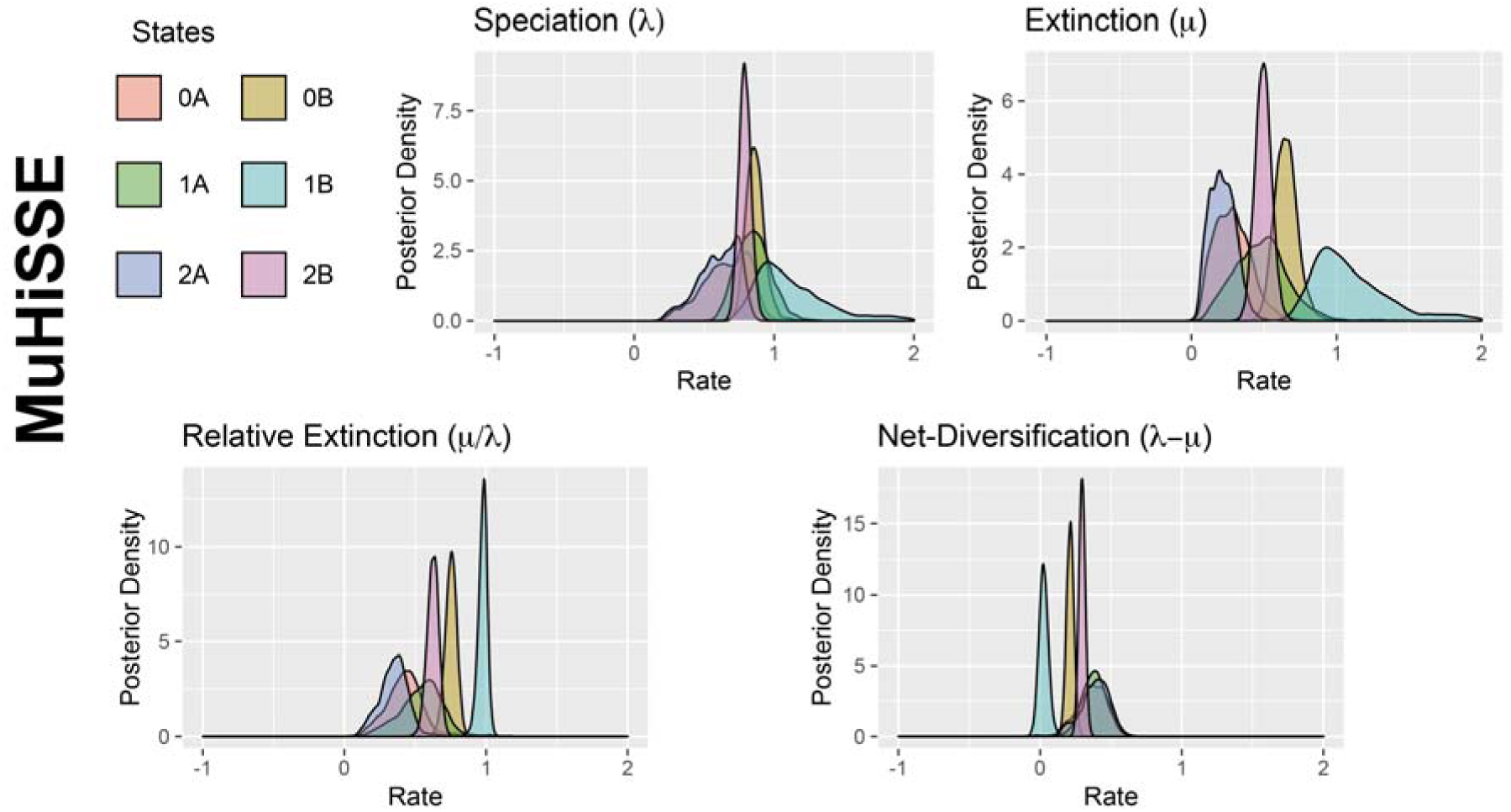
Trait-dependent posterior distributions of diversification rates estimated through MuHiSSE. Posterior densities of speciation (λ), extinction (μ), net-diversification (λ−μ) rates and relative extinction (μ/λ). Colors correspond to the posterior probabilities for a given state; note that differences in posterior distributions are larger between observed states (0, 1, 2) than between hidden states (A, B).

All SSE analyses ran in this study reached convergence, with traceplots showing adequate mixing and ESS values larger than 200 for all parameters (for many, this value was > 1000). The only exception was MuHiSSE, where ESS values were < 100 or 200 for many of the speciation and extinction parameters, even after combining the two runs, suggesting a possible lack of power. For this reason, the MuHiSSE model was excluded from model comparison using Bayes Factor power posteriors. The results from the Bayes Factor power posteriors, i.e. the marginal likelihood estimation for each model, are shown in Table 2; stepping-stone and path-sampling gave very similar values. The best-fitting model was HiSSE (ps = -16904.29), followed by MuSSE (ps=-16925.32) and BiSSE (ps = -16950.15). The CID2 model showed a worse fit to the data than any of the other SSE models (ps = -16956.41).

**Table 2.** Log-marginal likelihood values estimated with the Path Sampling (PS) and Stepping-stone (SS) methods for each trait-dependent diversification model applied in this study.

| SSE MODEL | Marginal Likelihood<br>SS | Marginal Likelihood<br>PS |
| --- | --- | --- |
| **BiSSE** | -16950 | -16950.15 |
| **HiSSE** | -16904.28 | -16904.29 |
| **MuSSE** | -16925.53 | -16925.32 |
| **CID2** | -16956.29 | -16956.41 |

## DISCUSSION

### Life Strategy Evolution in Blister Beetles

Our phylogenomic hypothesis supports previous works that considered parasitizing bees as the primitive life strategy of the hypermetamorphic clade of Meloidae (Bologna & Pinto 2001; Bologna et al. 2008). The sister-taxon relationship between the reciprocally monophyletic Meloinae and Nemognathinae suggests that the origin of this strategy dates back to the Eocene (Fig. 4). This general strategy has been retained by all but two of the descendent tribes, the non-sister taxa Mylabrini and Epicautini, which were able to experiment two separated non-simultaneous – homoplastic – host jump events from bees to grasshopper eggs (Fig. 4). Divergence time estimation and stochastic character mapping (Figs. 4, 5; Figs. S13, S14) indicate that the first host jumping event occurred in the Early Miocene along the stem-branch of Mylabrini, while the second, took place ten million years later, in the Mid-Miocene, along the stem-branch of Epicautini. Phylogenetic evidence for the independence of the two host jumping events is also provided by previous phylogenies (Bologna & Pinto 2001; Bologna et al. 2008).

Though there is ample evidence that host changes are common in nature (Sorenson et al. 2003; Wolfe et al. 2007; Giraud et al. 2010; Johnson et al. 2011; Forbes et al. 2017; Zhang et al. 2020), “dramatic” host-jumps, i.e. jumps to a new host species in a different family, order or phyla, are evolutionarily “rare”, compared to host shifts towards species closely related to the original host (Engelstädter & Fortuna 2019; Foster 2019; Braga et al. 2020). This is because establishing a sustainable relationship with a new host species represents an important challenge for parasites, which might require new morphological or physiological adaptations (Engelstädter & Fortuna 2019). Although the precise mechanism that facilitates a parasite jumping from one host to another is often unknown (Eppinger et al. 2006; Lowder et al. 2009; Shi et al. 2014), jumping probability success seems limited by the phylogenetic distance between the original and the new host (Engelstädter & Fortuna 2019; Foster 2019; Braga et al. 2020).

Host change in Mylabrini and Epicautini represents an example of “dramatic” host-jump: a shift between insect orders, from parasitizing hymenopteran larvae to feeding on orthopteran eggs. Hymenoptera and Orthoptera are phylogenetically distant, with their respective ancestors separated by more than 350 Ma (Misof et al. 2014; Song et al. 2015). Additionally, grasshoppers and bees exhibit very different development strategies, and their eggs differ markedly in chemical composition and properties (Hilker & Meiners 2008).

Grasshoppers of the family Acrididae are a dominant component of biodiversity in grassland ecosystems (Baldi & Kisbenedek 1997; Badenhausser et al. 2009), where species of Mylabrini and Epicautini are also abundant (Pinto 1991; Pinto & Bologna 1999; Bologna & Pinto 2002). Given that hosts and parasitoids share the same biome, and that food resource is abundant (i.e. grasshopper egg-pods), one could envisage continuous attempts by the parasitoid to shift to a new host, until a successful jump was achieved, not once, but twice independently over the evolutionary history of Meloinae. Other equally abundant insects in this biome are beetles of the families Tenebrionidae and Chrysomelidae.

However, the highly complex hypermetamorphic life cycle of Meloidae imposes severe evolutionary constraints, limiting a potential shift towards a non-orthopteran or hymenopteran host. A regular life cycle of a species in Nemognathinae and Meloinae spans a minimum of one year (Figs. 1, 2), but some have been recorded up to six years (Horsfall 1941; MacSwain 1956; Selander & Mathieu 1964; Selander & Weddle 1969). Stable temperature and soil moisture are the main factors governing the survival of the different larval stages (Erickson & Werner 1974a; Zhu et al. 2006). As in the case of bee nests, acridid egg-pods meet the requirements of a stable environment with constant temperature and soil humidity in which the successive larval types of the blister beetle may complete their development. This “ecological match” between the old and the new host, seems to be a common requirement for host-jump in other parasitoid lineages, such as velvet ants (Hymenoptera: Mutillidae) that were able to shift hosts from Hymenoptera to ant-nest dwelling Coleoptera with enclosed larvae (Chrysomelidae: Clythrini) (Brothers et al. 2000). Intriguingly, there are reports of larvae of *Cyaneolytta* (Meloinae) being phoretic on Carabidae, although their feeding habits are unknown (Di Giulio et al. 2003).

But, host-jump is not the only relevant change that occurred along the evolution of hypermetamorphic Meloidae. A change in the way first instar larvae reaches the bee-host from active crawling on the ground, to phoresy, occurred multiple times on the history of Meloidae (Fig. 7, Fig. S15). This inference agrees well with Bologna & Pinto (2001) and Bologna et al. (2008), who separately pointed out that bee-parasitism and non-phoresy were the ancestral states of hypermetamorphic Meloidae.

The homoplastic nature of phoresy in Meloidae translates in that traditionally regarded as natural tribes on the basis of morphology, such as Lyttini (non-phoretic), or Meloini (phoretic), become non-monophyletic in our mitogenomic phylogeny, including now genera with different life-history strategies. Lyttini, as currently defined, is composed of three non-sister lineages at least (Fig. 4): one lineage (Lyttini I) formed by *Lytta* (*L*. *caraganae*) and *Oenas* (*O*. *fusicornis*); a second lineage (Lyttini II) formed by *Berbermeloe* (*B. payoyo*), and a third (Lyttini III) lineage represented by *Lagorina sericea*. Bologna et al. (2008) already questioned the monophyly of Lyttini, and considered this tribe as a “non-phoretic taxonomic entity” where lineages with phylogenetic uncertainty were placed.

Unlike Lyttini, Meloini remains monophyletic if the genus *Spastonyx* is excluded, but again, Meloini is not characterized by a common life-history strategy: *Physomeloe* is a non-phoretic bee-parasitoid, which inclusion in Meloini, supported by our phylogenetic hypothesis (Fig. 4), was already suggested by Bologna et al. (2008). The phylogenetic position of the phoretic *Spastonyx* has long been a mystery. This genus comprises only two small-sized species that inhabit the arid areas of northern Mexico and southern United States (López-Estrada et al. 2018). Our phylogeny suggests that *Spastonyx* is more closely related to Pyrotini (non-phoretic) than to the rest of Meloini or Lyttini, where it has been doubtfully included (Pinto & Selander 1970; Bologna & Di Giulio 2011).

Bologna et al. (2008) suggested that phoresy was evolutionarily advantageous in Meloidae because it enhances the ability of the first instar larvae to reach the host, thus ensuring the food resource availability. Actively latching to the host, rather than wandering around to locate the nest, can also be seen as a strategy to save energy (Baumann et al., 2018). In other organisms, phoresy acts as a dispersal strategy that ensures genetic exchange between populations and homogenizes them (Athias-Binche et al. 1993; DiBlasi et al. 2018; Opatova & Št’áhlavský 2018). Conversely, the ancestral “bee-by-crawling” strategy would not be advantageous in a similar scenario. The first instar larva of the hypermetamorphic Meloidae is highly mobile, but unlikely able to cover great distances (Erickson & Werner 1974a; Selander & Weddle 1969). Activity periods, host searching behavior and longevity of these larvae are highly influenced by the environmental temperatures and soil moisture (Erickson & Werner 1974b). Without a clear preference for oviposition site (Erickson & Werner 1974a; Selander & Weddle 1969), with limited dispersal capabilities and under adverse climatic conditions, first instar larvae are forced to reach the host as quickly as possible. Under this scenario, a “wandering”, “bee-by-crawling” strategy seems the least suitable.

Some studies, especially in mites (Brown & Wilson 1992; Athias-Binche et al. 1993), suggest that phoresy may also induce speciation by “host-phoretic specialization”, in which the parasite modifies certain traits to ensure successful latching to a specific host.

Some lineages such as *Meloe* exhibit large morphological diversity in larval traits related to phoresy (Bologna 1983; 1991; Bologna & Pinto 2001), including variable development of a pygopod to crawl on vertical surfaces, or variable degree of abdominal sclerotization (Cros 1940; MacSwain 1956; Kaszab 1969; Selander 1964; Pinto & Selander 1970; Bologna & Pinto 2001; Bologna et al. 2008). Meeting thus, some of the evolutionary requirements involved in a host-phoretic specialization process.

### Host-Jump and Phoresy Triggered Diversification in Meloidae

Independent host jumping events seem to have triggered an increase in the rate of diversification in Meloinae. BiSSE and MuSSE indicate that the grasshopper specialist tribes Mylabrini and Epicautini, which are the most speciose, exhibit higher speciation rates than those of their sister taxa, non-phoretic, bee-parasitoid lineages (Figs. 5, 7, Fig. S16).

Evidence that host specialization can be a powerful driver of diversification comes from many different organisms, including phytophagous insects (Ehrlich & Raven 1964; Forbes et al. 2017), pathogenic fungi (Giraud et al. 2010), helminth worms (Zietara & Lumme 2002), avian brood parasites (Sorenson et al. 2003), and ectoparasitic arthropods (Johnson et al. 2011) among others. Adaptation to the new habitat (host) can induce reproductive barriers in a relatively small number of generations (Hendry et al. 2007), and thus accelerate the rate of speciation events, sometimes leading to patterns consistent with adaptive radiations (Zietara & Lumme 2002; Farrel & Sequeira 2004; Fordyce 2010; Karvonen & Seehausen 2012; Forbes et al. 2017; Bush et al. 2019).

Host specialization is often associated with major changes in morphology, physiology, anatomy, or reproductive mechanisms (Barnhart et al. 2008; Schulte et al. 2010; Turrisi & Vilhelmsen 2010). But, so far, no set of traits shared between the larvae of grasshopper specialists Epicautini and Mylabrini has been found (Bologna & Pinto 2001; Bologna et al. 2008). This suggests that no prior morphological adaptations or key innovations were involved in the evolution of the grasshopper specialization strategy. This is in accordance with the concept of “ecological fitting” (Janzen 1985), in which the formation of a new interaction does not require the evolution of new traits; instead, it is based on traits developed for previous host-parasite interactions and “co-opted” for a new interaction given the right conditions (Agosta 2006).

In addition to host-jump, phoresy is another driver of diversification in hypermetamorphic blister beetles. Our SSE analyses indicate that phoretic bee-parasitoid lineages exhibit higher net diversification rates and lower relative extinction rates than non-phoretic bee-parasitoid lineages, being the latter the ancestral condition (Figs. 6-8; Fig. S15). Although phoresy is a diversification driver, it might be acting either through host-phoretic specificity, or as a strategy to ensure panmixia and food availability (Opatova & Št’áhlavský 2018), however, these alternative scenarios have not been statistically tested. Our SSE analyses demonstrate that phoresy is a life strategy as effective as a change of host to increase diversification rates in hypermetamorphic Meloidae. In fact, no significant differences in net diversification or relative extinction rates were found between phoretic bee-parasitoids and non-phoretic grasshopper specialists (Figs. 7, S16).

### Historical Contingency and Evolutionary Dead Ends in Hypermetamorphic Meloidae

While trying to understand why certain groups diversify more than others, the idea of a single key factor promoting elevated diversification rates (e.g., the evolution of a morphological innovation, the invasion of a new isolated environment, or the effect of a mass extinction event depleting extant diversity) has been a dominant one in the literature (Hodges & Arnold 1995; Hunter & Jernvall 1995; De Queiroz 2002; Donoghue 2005). Yet, despite the numerous recent studies testing for an association between trait evolution and diversification, few of them have found evidence of a single trait driving a shift in diversification rates (Lagomarsino et al. 2017; Condamine et al. 2018; Moharrek et al. 2019). Instead, the dominant pattern is one in which bursts of diversification are explained by the confluence of multiple factors, sometimes acting at unison (Donoghue & Sanderson 2015), sometimes in a sequence (Donoghue 2005), or contingent upon one another (Givnish et al. 2015).

Our study of diversification in blister beetles supports this idea of multiple interacting factors. Although phoresy and host-jump may both induce speciation by host specialization, neither host-type nor phoresy themselves can explain the pronounced differences in species richness observed among hypermetamorphic Meloidae tribes and genera. Instead, it is the parallel innovation brought by these two traits that directly impacts the diversification dynamics of Nemonagthinae and Meloinae. Though BiSSE detected a signal of trait-diversification dependency with host-type (Fig. 5), HiSSE and MuSSE (Figs. 6, 7) pointed out that host-type alone cannot explain the observed differences in species richness; these results were robust against the introduction of hidden factors (Fig. 8) or phylogenetic uncertainty (Fig. S16).

On the other hand, these two traits have not acted simultaneously or synergistically as “synnovations” (Donoghue 2005; Donoghue & Sanderson 2015). Non-phoretic lineages can be either bee-parasitoids or grasshopper specialists, but all phoretic lineages are bee-parasitoids (i.e. there are not phoretic grasshopper specialists). The fact that adult grasshoppers never return to the egg-pods once the oviposition takes place, makes phoresy in grasshopper specialist lineages highly unlikely. In contrast, the social behavior of bees, characterized by long-term offspring rearing, makes phoresy a highly efficient strategy to ensure food resources for lineages that are bee parasitoids (Danforth 2007). Thus, in our analysis, we could not treat these two traits as independent binary characters and use an extended four-state SSE model to examine their joint evolution (Nakov et al. 2019).

Instead, a three-state MuSSE model (Fig. 6) was used to examine how phoresy interacts with host-type in explaining rate heterogeneity between bee and grasshopper parasitoids, as detected by BiSSE (Fig. 5). Our results show that phoresy and grasshopper specialization acted as two independent causal forces behind the diversification rate heterogeneity in blister beetles.

Parasitoidism on bees could be seen as a peculiar example of “historical contingency” in evolution (Gould 1989; Losos et al. 1998; Vermeij 2001; De Queiroz 2002). Beatty (2006) proposed a distinction between “causal contingency”: i.e. a change in one character that limits the evolutionary outcome of a different character, and “unpredictable contingency” (Gould 1989). In our study, parasitoidism on bees fits the definition of “causal contingency” (Beatty 2006; Losos et al. 1998): phoresy could not be achieved if hypermetamorphic Meloidae never acquired bee-parasitism as a life strategy. On the other hand, a host-jump towards feeding on orthopteran eggs would represent an example of “unpredictable contingency” (Gould 1989). In other words, the onset of the “bee-by-crawling parasitoidism” in the hypermetamorphic Meloidae acted as a “historical constraint” in two different ways: as a causal factor enabling further specialization (phoresy) or by leaving an open window for unpredictability, allowing for a dramatic innovation (a host-jump) along non-sister lineages.

These two “derived” life strategies in Meloinae, phoresy and grasshopper specialization, could represent adaptive radiation drivers (Erwin 1992; Gavrilets & Losos 2009; Givnish et al. 2014; Simões et al. 2016) – seen as an “ecological opportunity” (i.e. the adaptation to a new environment such as the grasshopper egg-pods in Mylabrini and Epicautin), or as a “key innovation” (the evolution of a phoretic strategy in bee-parasitoid lineages). On the contrary, the life-history strategy of the ancestors of Meloinae and Nemonagthinae is associated with a lower net diversification rate and a significantly higher relative extinction rate than the “bee-by-phoresy” or “grasshoppers-by-crawling” strategies. Our MuSSE reconstruction (Fig. 7) indicates that transitions to the latter strategies occurred at least five times independently over the evolution of the hypermetamorphic Meloidae, always accompanied by an increase in diversification rate. Under these circumstances the “bee-by-crawling” or “wandering” strategy could be tending towards an “evolutionary dead end”. Interestingly, stochastic character mapping (Fig. S15b) suggests that the “bee-by-crawling” strategy might have been ancestrally present in *Cissites*, a species-poor genus subtended by a long stem-branch, belonging to the species-poor tribe Horiini (15 species; Pinto & Bologna 1999; Bologna & Pinto 2002), which is the sister-group of the more speciose tribe Nemognathini (Table 2). It is possible that the “bee-by-crawling” strategy was initially present within Nemognathinae, but because of the high extinction rate associated with this condition, no lineage exhibiting it presently survived in the subfamily. “Bee-by-crawling” lineages within Meloinae might also suffer a similar fate. This supports the idea that the non-phoretic bee-parasitoid lineages of Meloidae act as “depauperons”, the “flip side” of evolutionary radiations, species-poor clades with old divergence times that could be slowly deriving towards an evolutionary dead end (Donoghue & Sanderson 2015). In other words, the host-jump to grasshopper eggs and the acquisition of the phoretic behavior in bee parasitoids could be evolutionary pathways by which the ancestral hypermetamorphic Meloidae lineages “escaped” extinction.

The rushing forward of blister beetles to escape possible extinction determined by their ancestral complex life strategy is remarkable. The low net levels of diversification for the bee-by-crawling strategy suggest that lineages retaining the ancestral condition (bee-by-crawling parasitoids) would be slowly disappearing until extinction through progressive depauperation (Donoghue & Sanderson 2015). However, the ability of blister beetles to explore new evolutionary scenarios (López-Estrada et al. 2019) seems to constantly open new pathways to escape extinction (phoresy, host-jumps). It does not seem likely that selective pressures driving homoplasy acted in two characters (host-jump and phoresy) that do not set the conditions for the success of the other, unless blister beetles were constantly exploring alternative life strategies, breaking the evolutionary restrictions imposed by their phylogenetic history. In this line, López-Estrada et al. (2019), suggested that the facility to explore different major axis of change (in morphological body ground plan) allowed a tribe of Meloidae to escape climatic extinction later in the evolution of the group. Following Vermeij (2006) reasoning, macroevolutionary change associated to life-history strategies in Meloidae is largely an effect of contingency, but since host-jump and phoresy occurred multiple times following speciation events (therefore somewhat predictable), replicate independent ecological analysis at an accessible (e.g. recent) temporal scale, are attainable. This situation opens a window to explore at a microevolutionary scale the mechanisms that originate large-scale macroevolutionary change.

Hidden-SSE models are typically used to discard Type I error or “false positives” when examining a causal association between trait evolution and diversification-rate heterogeneity (Condamine et al. 2018; Fernandez et al. 2018; Gajdzik et al. 2019; Nakov et al. 2019). Little attention is paid to negative results (Donoghue, 2005), for example when the HiSSE model does not support the association between the focal trait and increased diversification rates (Fig. 6). In our study, we showed that HiSSE may be useful to identify the actual “hidden” trait that is driving diversification dynamics in the group. In hypermetamorphic blister beetles, rather than host-jump, as originally hypothesized, diversification rate heterogeneity is the product of changes in life strategy, either by developing phoresy to secure the resource (food) or by a dramatic host-jump to a different order (Orthoptera).

## SUPPLEMENTARY MATERIAL

The following supplementary figures can be found within the online supplementary material.

**Figure S1.** The backbone ultrametric mitogenomic tree, obtained using BEAST using the AA+rNT-matrix showing the location of nodes for which we simulated subtrees with a number of taxa equal to the current taxonomic species richness (right inset). Subtrees were simulated under an integrative approach (see main text) using functions from *TreePar*, *geiger*, *TESS* and ape R packages.

**Figures S2-S6.** Directional Acyclic Graphs (DAGs) showing parameter dependency and priors used in the five hierarchical Bayesian Diversification models examined here: S2: BiSSE, S3: HiSSE, S4: MuSSE, S5: MuHiSSE, S6: CD2. More details on models can be found in the main text of the article. DAGs were plotted using GraphViz.

**Figure S7.** Gene arrangement and general structure of the mitogenome of hypermetamorphic Meloidae (subfamilies Nemognathinae and Meloinae).

**Figure S8.** Phylogenetic hypothesis obtained using Maximum Likelihood Estimate (MLE) with RAXML on (A) NT and (B) AA+rbNT matrices. Bootstrap support values are shown above nodes.

**Figure S9**. Bayesian Majority-rule consensus tree obtained with MrBayes using (A) NT and (B) AA+rbNT matrices. Posterior probability (pp) clade support values are shown above nodes (∼88% of the nodes are supported with a pp value equal to 1; see Fig. 3 for additional information).

**Figure S10.** Bayesian Majority-rule consensus tree obtained with PhyloBayes under the CT-Poisson model using (A) NT and (B) AA matrices. Posterior probability clade support values are shown above nodes.

**Figure S11.** Lineage divergence times as estimated in BEAST using Bayesian relaxed clocks using the AA+rbNT matrix. Mean ages and 95% High Posterior Density (HPD) values are shown near to each node; violet horizontal bars represent the range of the HPD. **Figure S12**. Plot of pairwise differences of speciation (A) and extinction (B) rates between states 0 and 1 estimated by BiSSE. The histogram shows the distribution of differences across the MCMC posterior distribution; the red line indicates the 0 value (no differences); black dotted lines indicate the confidence intervals.

**Figure S13.** Results from the BiSSE model. (A) Maximum A Posteriori (MAP) tree showing reconstructed ancestral states, indicated with colors; size of circles represents marginal posterior probabilities. (B) MAP tree showing the number and timing of transition events between states reconstructed along branches using stochastic character mapping; inset to the left indicates marginal posterior probability for the inferences depicted as colors.

**Figure S14.** Results from the HiSSE model. (A) Maximum A Posteriori (MAP) tree showing reconstructed ancestral states, indicated with colors; size of circles represents marginal posterior probabilities. (B) MAP tree showing the number and timing of transition events between states reconstructed along branches using stochastic character mapping; inset to the left indicates marginal posterior probability for the inferences depicted as colors.

**Figure S15.** Results from the MuSSE model. (A) Maximum A Posteriori (MAP) tree showing reconstructed ancestral states, indicated with colors; size of circles represents marginal posterior probabilities. (B) MAP tree showing the number and timing of transition events between states reconstructed along branches using stochastic character mapping; inset to the left indicates marginal posterior probability for the inferences depicted as colors.

**Figure S16.** Phylogenetic uncertainty analysis. Plot of pairwise differences of diversification rates among states (0, 1, 2) estimated by the MuSSE model. The histogram represents the distribution of these values across the 100 backbone+simulated subtrees phylogenies. The distribution is centered in 0 (represented by the red line) for states 1 and 2 (i.e. there are no differences in diversification rates across the simulated phylogenies). However, there are significant differences for the other two pairwise comparisons, confirming the results reported in Figure 6.

**Table S1.** List of specimens included in this study, locality data, ID number of the Entomological Collection of the Museo Nacional de Ciencias Naturales (MNCN-CSIC) and GenBank (GB) accession numbers for the mitochondrial (mt) genomes.

## AUTHOR CONTRIBUTIONS

EKLE, IS and MGP conceived the idea. EKLE and MGP carried out the fieldwork. JEU and SA generated the molecular data. EKLE, JEU and SA assembled and annotated the mitogenomes, and performed phylogenetic inference with help from IS. EKLE and IS designed and conducted the diversification analyses. EKLE wrote the manuscript together with IS and MGP, with contributions from SA and JEU.

## ACKNOWLEDGMENTS

We are thankful to Alberto Sánchez-Vialas, José Luis Ruiz, Ernesto Recuero, Jorge Gutiérrez-Rodríguez, Paloma Mas-Peinado, Nohemí Percino-Daniel, and Gonzalo García for their help with fieldwork; Yolanda Jiménez Ruiz for assistance in the laboratory, and Edna Gabriela López Estrada for help with R scripting. We especially thank Rogelio Delgado Román for generously providing us with additional computational resources to conduct this study. We thank the Escobar Biological Station (Cazalla de la Sierra, Spain) and all members of the Biodiversity Screening Laboratory for support during the COVID19 confinement.

EKLE is supported by a doctoral scholarship from CONACyT–Mexico (330519/472100). SA has been supported by the Spanish Ministry of Science, Industry and Innovation, (BES-2014-069575) and the SponGES H2020 grant (BG-01-2015.2, agreement number 679849-2). JEU was supported by Peter Buck Postdoctoral Fellowship Program from the Smithsonian Institution (2017–2019) and is currently supported by the Atracción Talento de la Comunidad de Madrid Fellowship Program (REFF 2019-T2/ AMB-13166).

