## supplementary material for "Diversification dynamics of hypermetamorphic blister beetles (Meloidae): Are homoplastic host shifts and phoresy key factors of a rushing forward strategy to escape extinction?"

**Figure S1.** The backbone mitogenomic tree, showing the location of nodes for which we simulated subtrees with a number of taxa equal to the current taxonomic species richness (right inset).

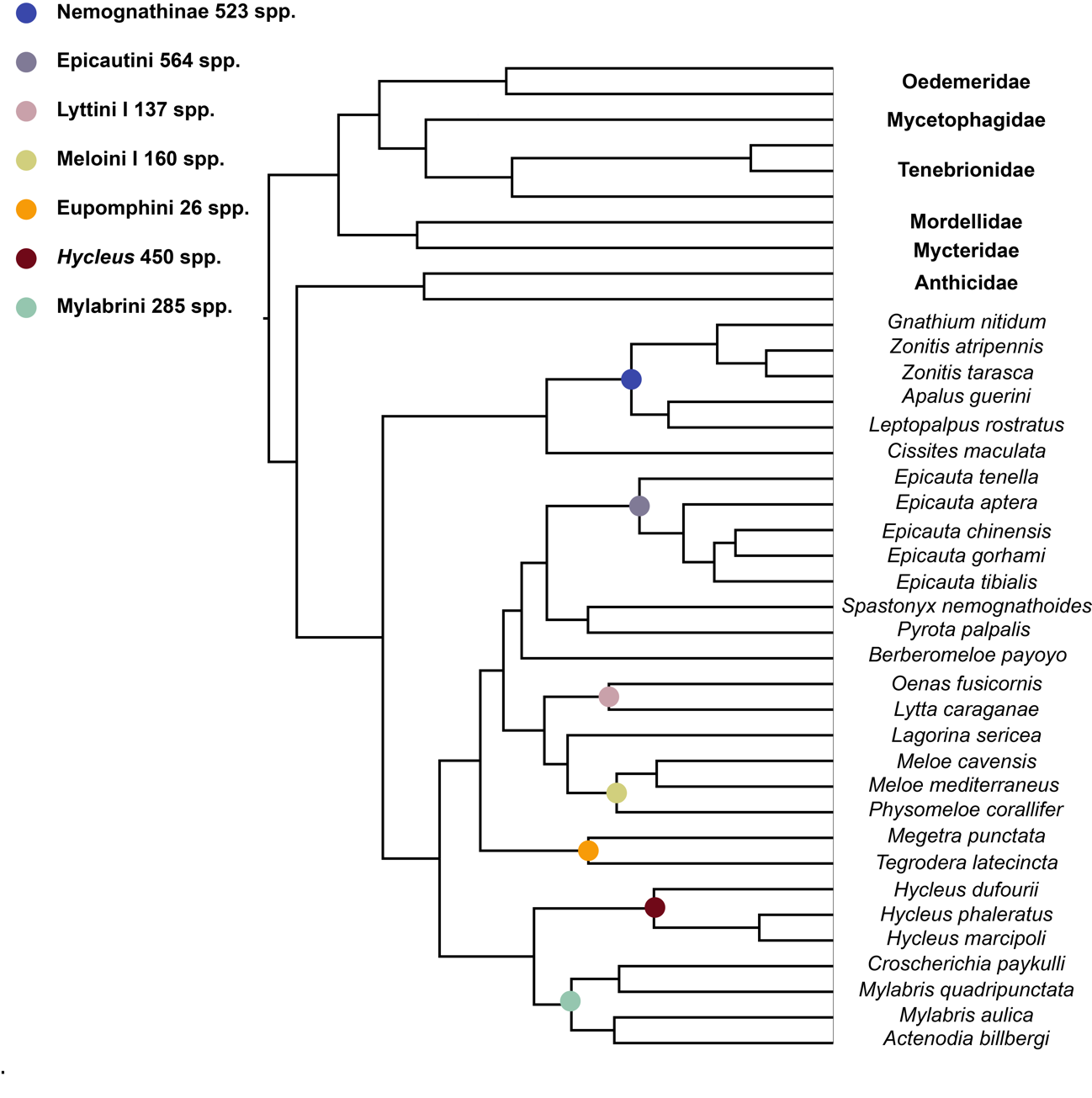

**Figure S2.** Directional Acyclical Graphs (DAGs) showing the parameter dependency and priors used in the hierarchical Bayesian Diversification model BiSSE.

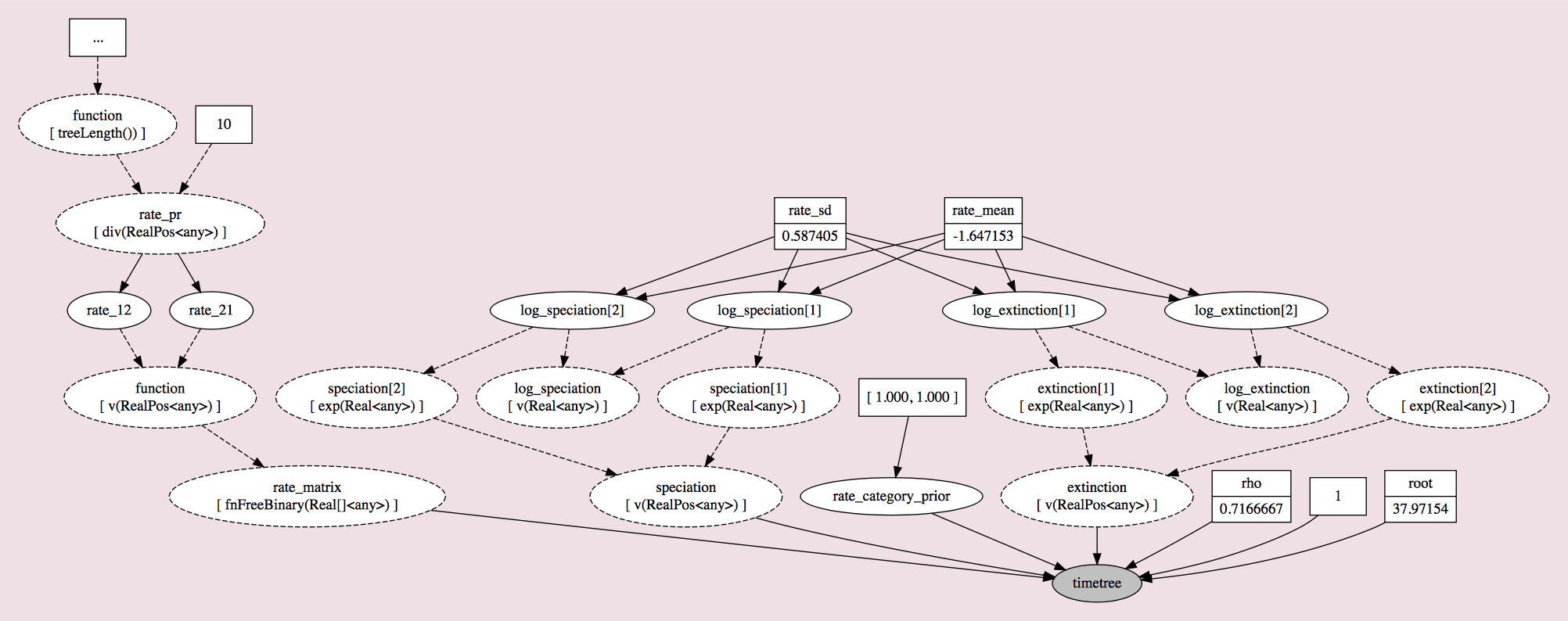

**Figure S3.** Directional Acyclical Graphs (DAGs) showing the parameter dependency and priors used in the hierarchical Bayesian Diversification model HiSSE.

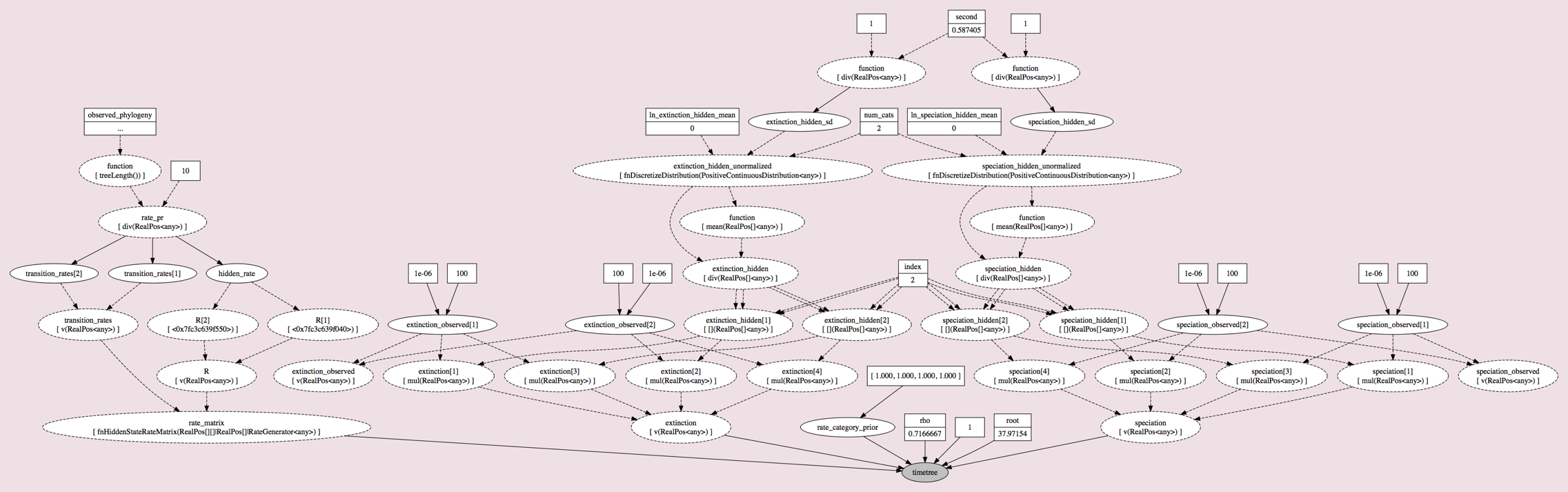

**Figure S4.** Directional Acyclical Graphs (DAGs) showing the parameter dependency and priors used in the hierarchical Bayesian Diversification model MuSSE.

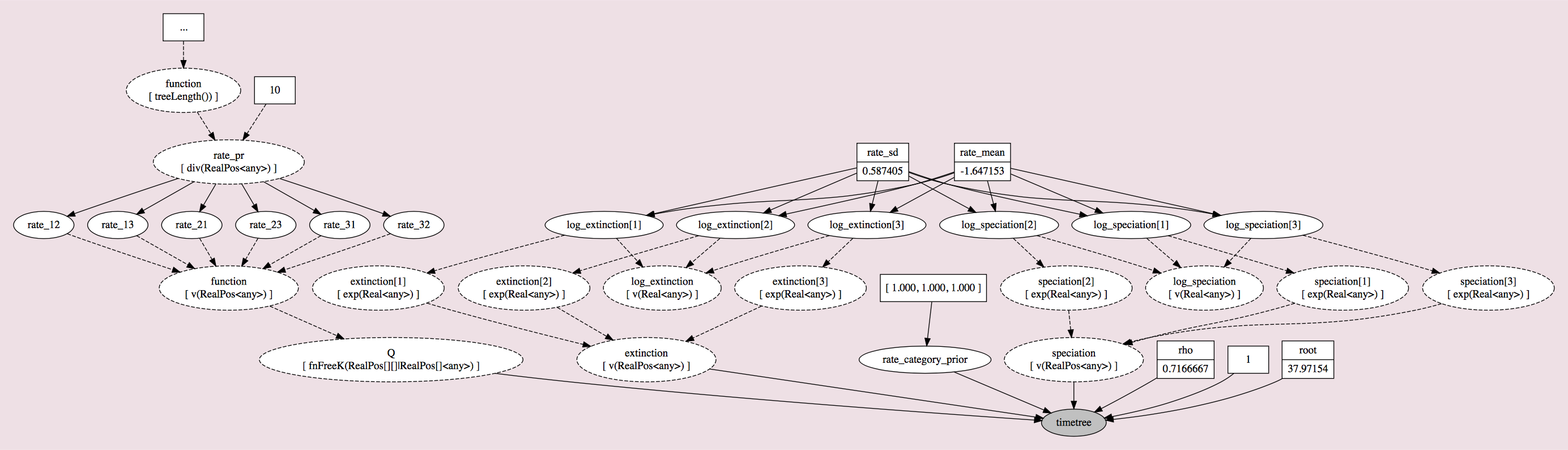

**Figure S5.** Directional Acyclical Graphs (DAGs) showing the parameter dependency and priors used in the hierarchical Bayesian Diversification model MuHiSSE.

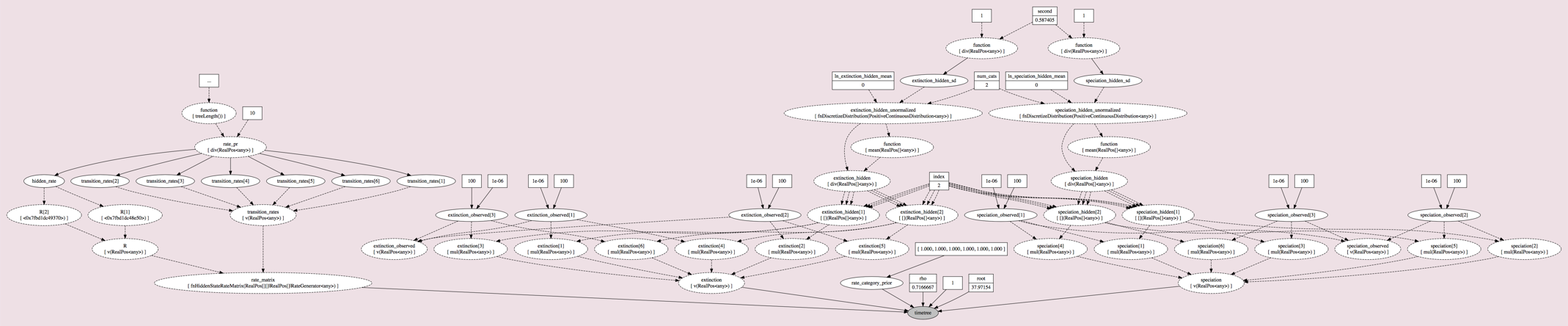

**Figure S6.** Directional Acyclical Graphs (DAGs) showing the parameter dependency and priors used in the hierarchical Bayesian Diversification model CID2.

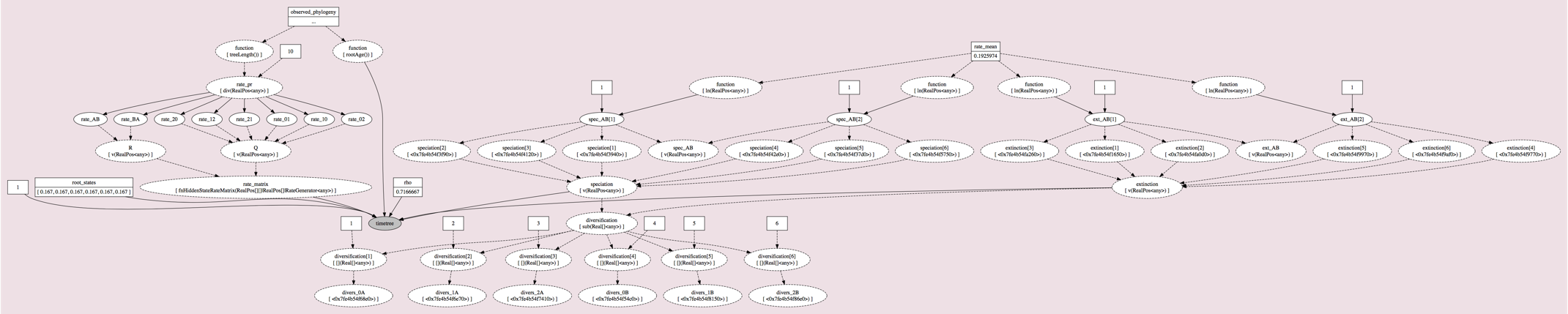

**Figure S7**. Gene arrangement and general structure of the mitogenome of the hypermetamorphic clade of Meloidae.

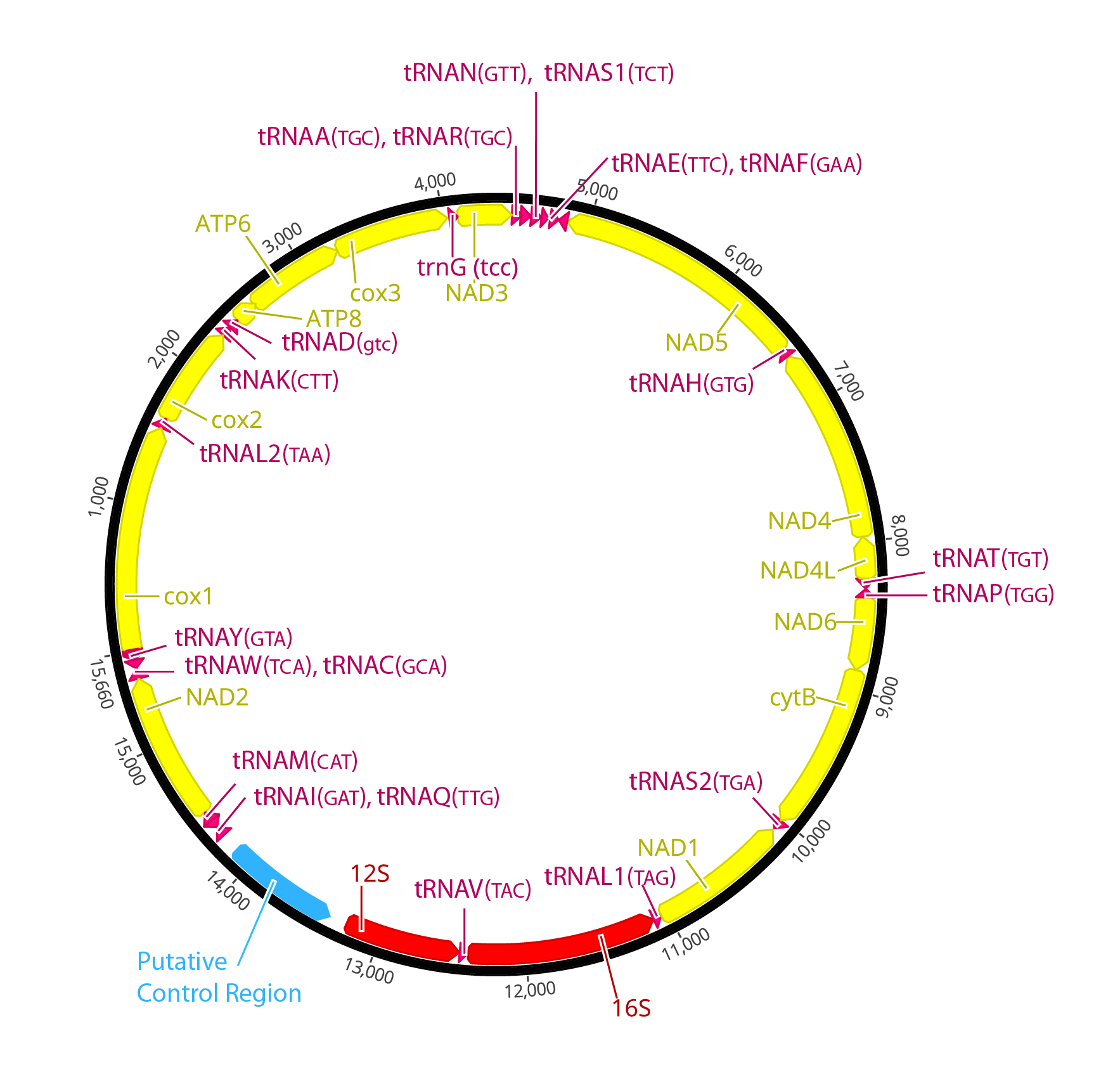

**Figure S8.** Maximum likelihood estimate (MLE) of the phylogeny obtained with RAXML using (A) NT-matrix and (B) AA+rbNT matrix. Bootstrap support values are shown above nodes.

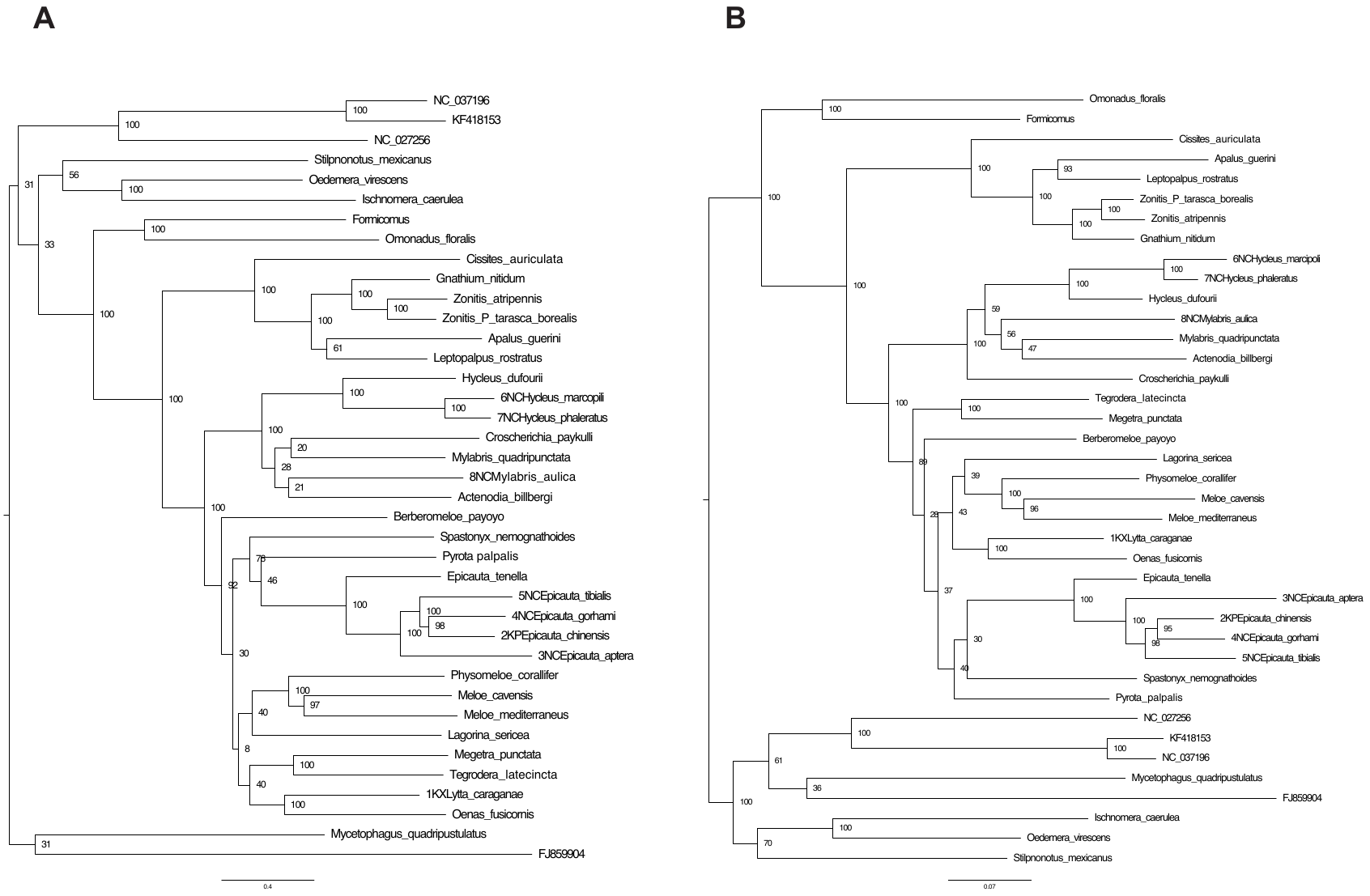

**Figure S9**. Bayesian Majority-rule consensus tree obtained with MrBayes using (A) NT-matrix and (B) AA+rbNT matrix. Posterior probability clade support values are shown above nodes.

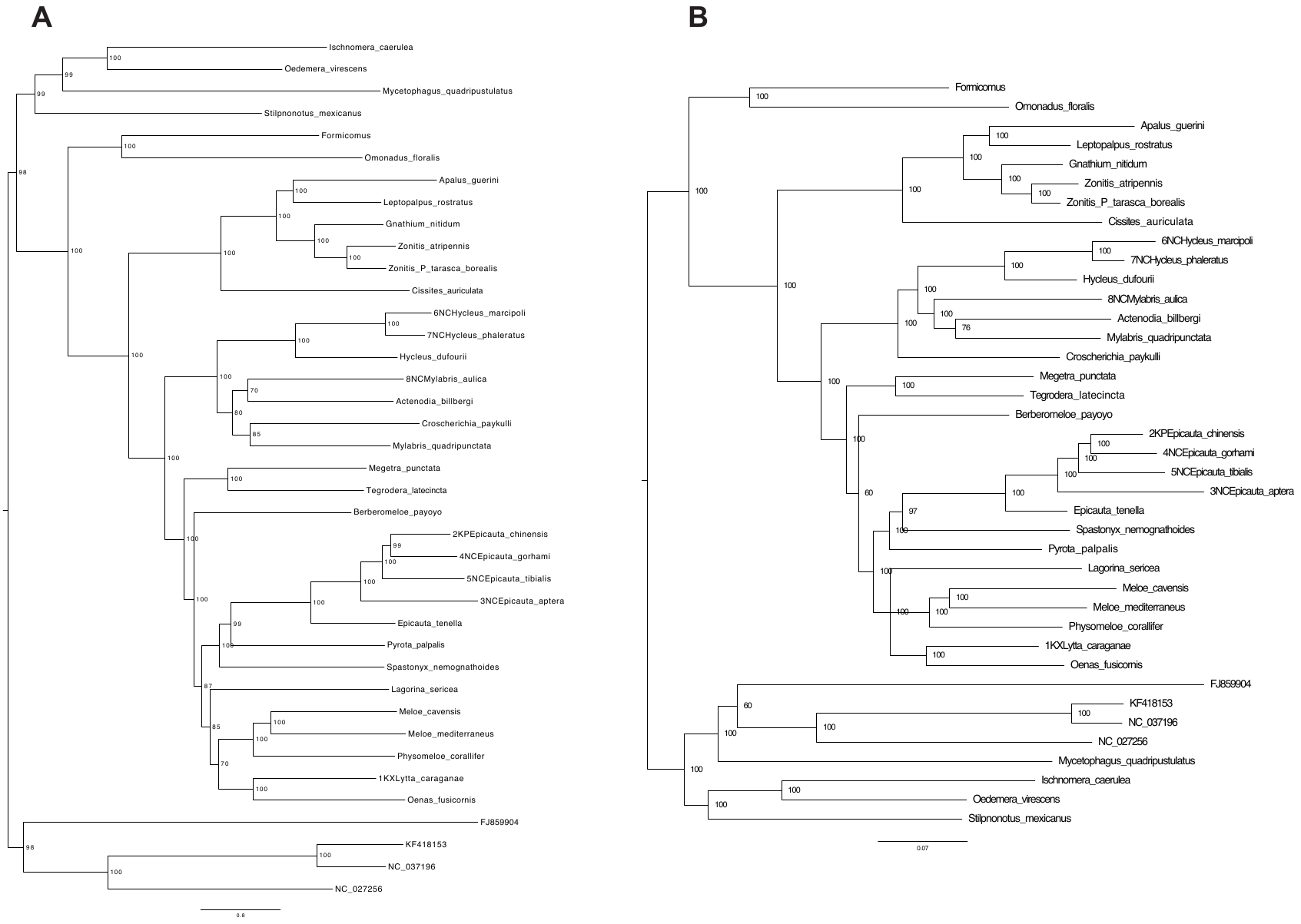

**Figure S10:** Bayesian Majority-rule consensus tree obtained with PhyoBayes under the CT-Poisson model using (A) NT-matrix and (B) AA matrix. Posterior probability clade support values are shown above nodes.

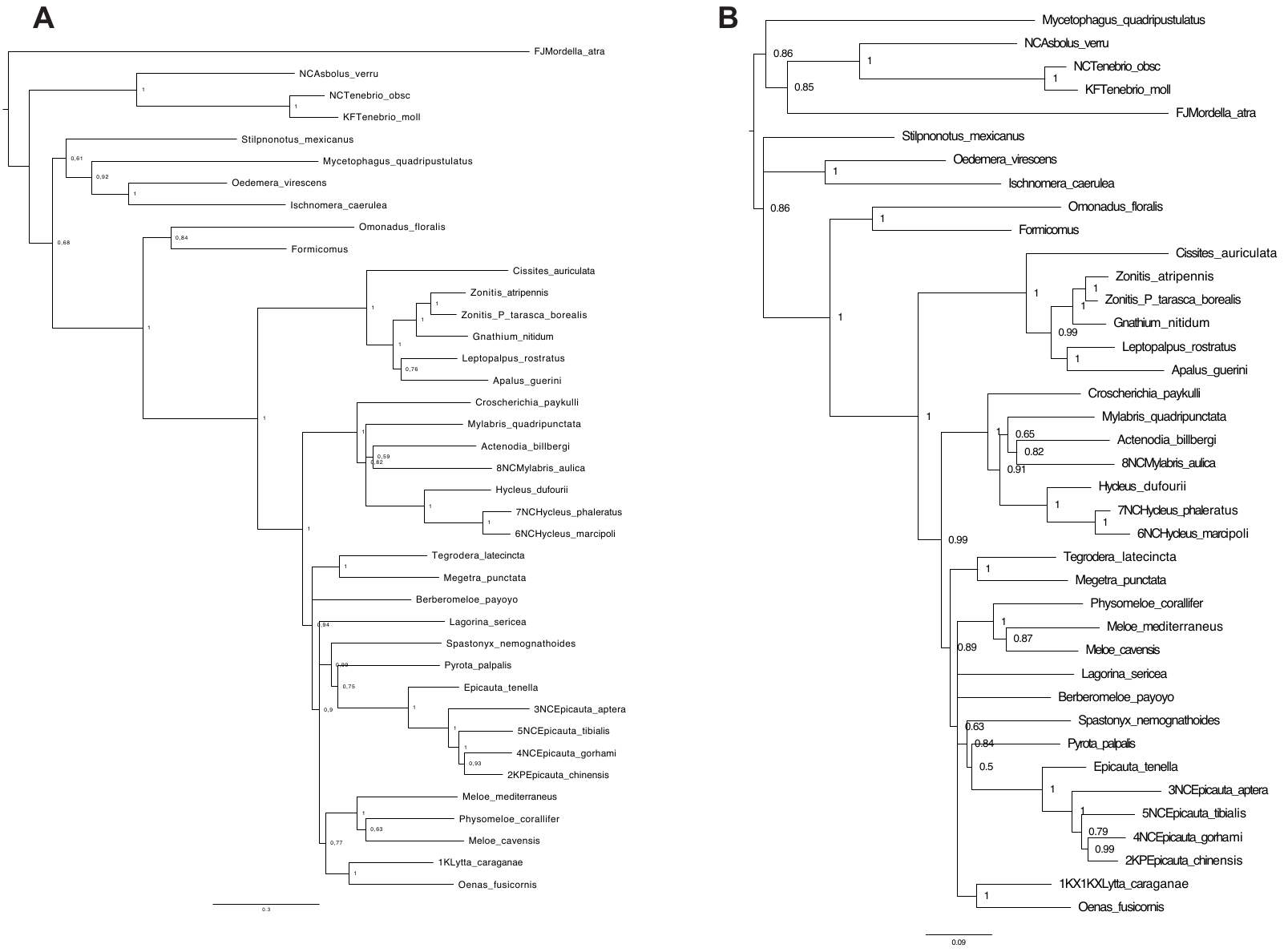

**Figure S11**. Lineage divergence times as estimated in BEAST using Bayesian relaxed clocks using the AA+rbNT matrix. Mean ages and 95% High Posterior Density (HPD) values are shown near to each node; violet horizontal bars represent the range of the HPD.

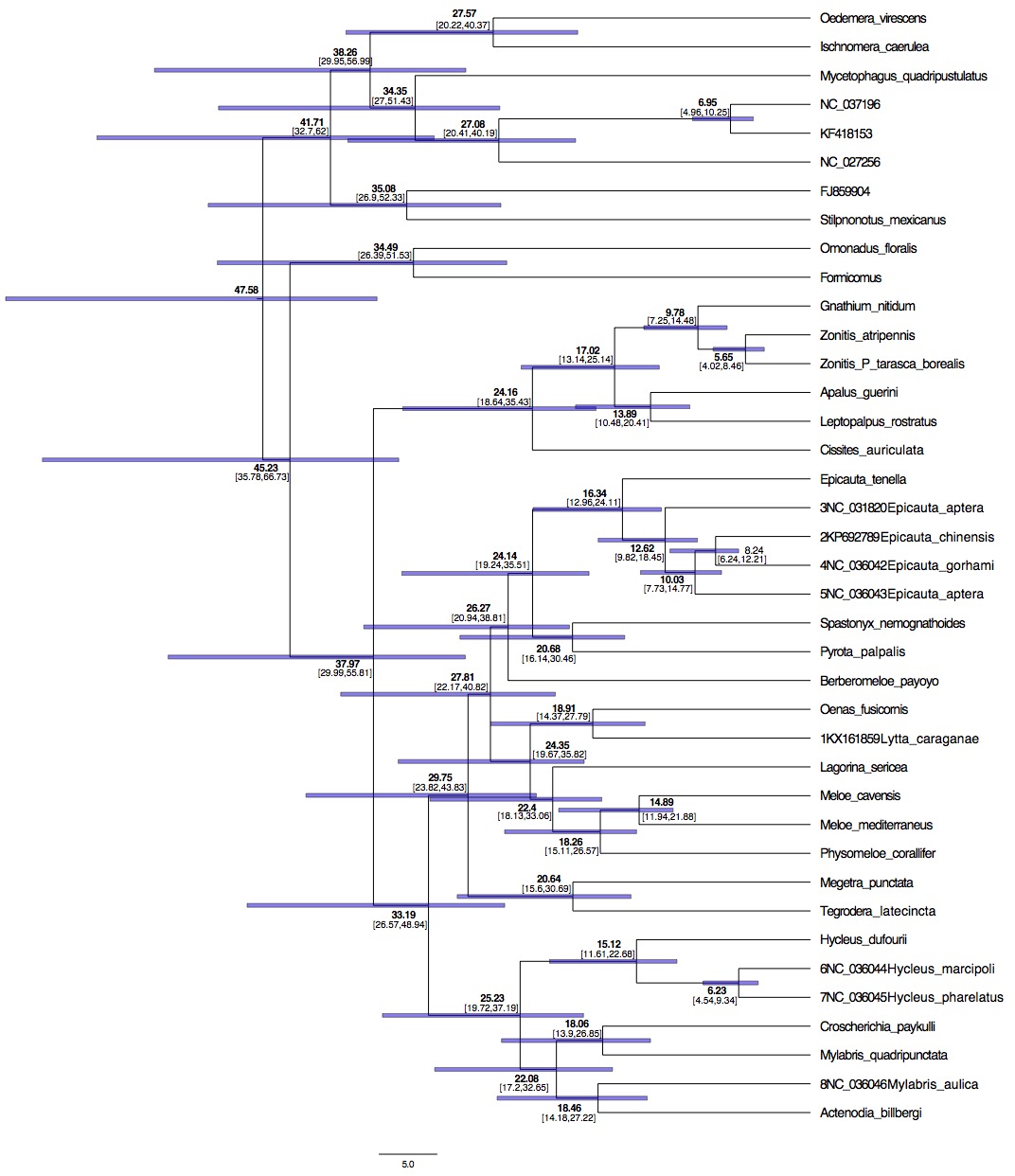

**Figure S12**: Plot of pairwise differences of speciation (A) and extinction (B) rates between states 0 and 1 estimated by BiSSE. The histogram shows the distribution of differences across the MCMC posterior distribution; the red line indicates the 0 value (no differences); black dotted lines indicate quartiles.

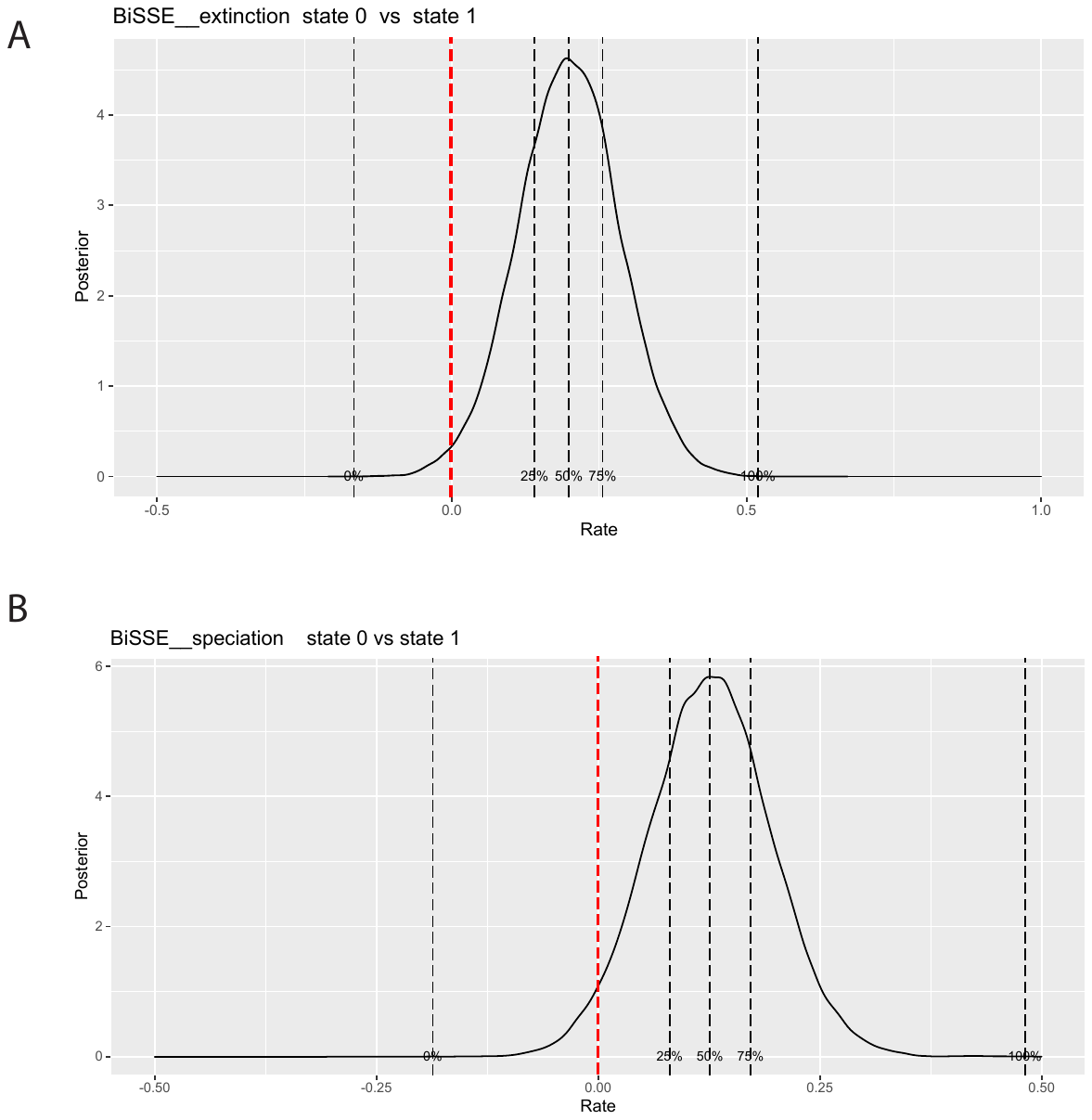

**Figure S13.** Results from the BiSSE model. (A) Maximum A Posteriori (MAP) tree showing reconstructed ancestral states, indicated with colors; size of circles represents marginal posterior probabilities. (B) MAP tree showing the number and timing of transition events between states reconstructed along branches using Stochastic Mapping; inset to the left indicates marginal posterior probability for the inferences depicted as colors.

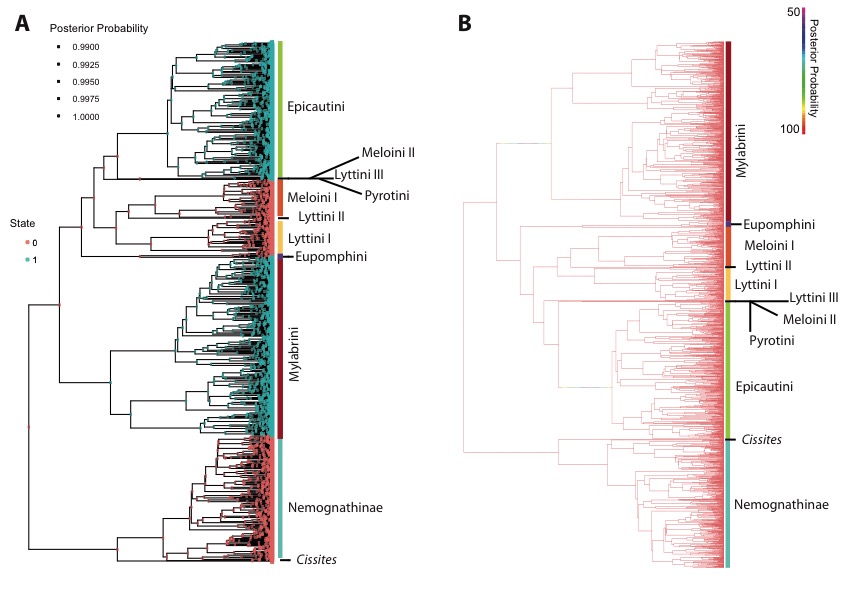

**Figure S14.** Results from the HiSSE model. (A) Maximum A Posteriori (MAP) tree showing reconstructed ancestral states, indicated with colors; size of circles represents marginal posterior probabilities. (B) MAP tree showing the number and timing of transition events between states reconstructed along branches using Stochastic Mapping; inset to the left indicates marginal posterior probability for the inferences depicted as colors.

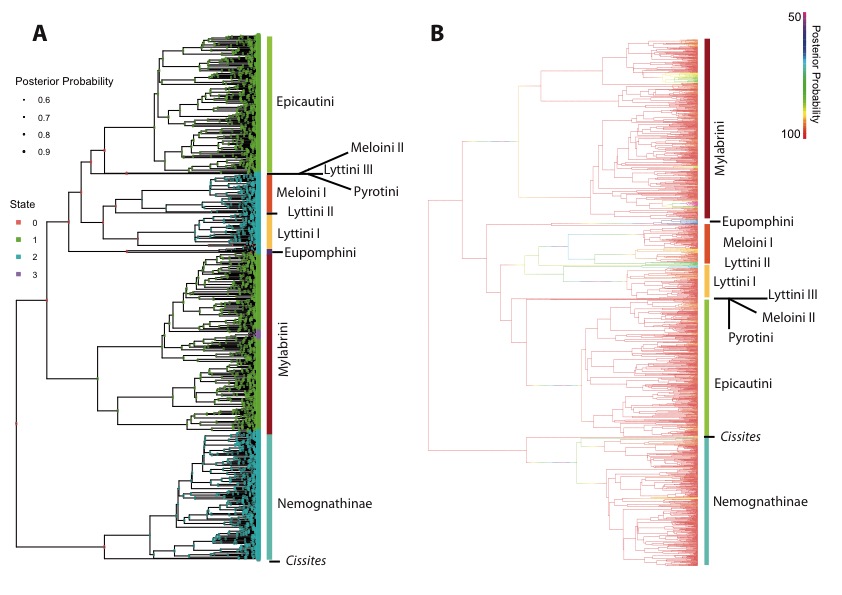

**Figure S15.** Results from the MuSSE model. (A) Maximum A Posteriori (MAP) tree showing reconstructed ancestral states, indicated with colors; size of circles represents marginal posterior probabilities. (B) MAP tree showing the number and timing of transition events between states reconstructed along branches using Stochastic Mapping; inset to the left indicates marginal posterior probability for the inferences depicted as colors.

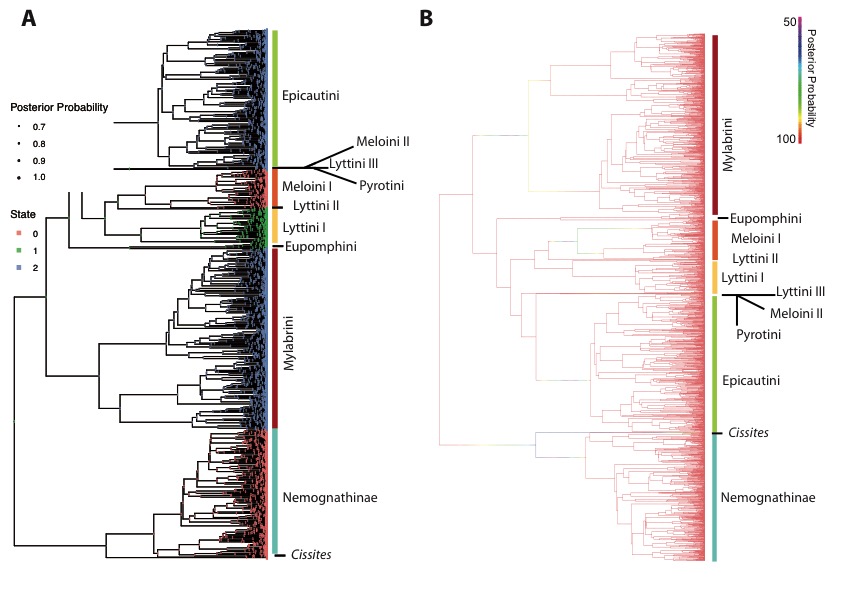

**Figure S16.** Phylogenetic uncertainty analysis. Plot of pairwise differences of diversification rates among states (0, 1, 2) estimated by the MuSSE model. The histogram represents the distribution of these values across the 100 backbone+simulated subtrees phylogenies. The distribution is centered in 0 (represented by the red line) for comparison between states 0 and 2 (i.e., there are no differences in diversification rates across the simulated phylogenies). However, there are significant differences for the other two pairwise comparisons (0 vs 1 and 1 vs 2), confirming the results reported in Figure 5.

**Table S1.** List of specimens included in this study, locality data, ID number of the Entomological Collection of the Museo Nacional de Ciencias Naturales (MNCN) and GenBank (GB) accession numbers for the mitochondrial (mt) genomes.

| **New mt genomes** | | | | | | | | | | | | | | | |
| --- | --- | --- | --- | --- | --- | --- | --- | --- | --- | --- | --- | --- | --- | --- | --- |
| **MNCN specimen number** | **Subfamily** | | **Tribe** | | **Species** | | | **Locality** | **Coverage** | | | **Length**  **bp** | | | **GB accession number** |
|  |  |  |  |  |  |  |  |  | # reads | Mean depth | |  | |  | |
| APP15148a | Meloinae | | Lyttini | | *Berberomeloe payoyo* Sánchez-Vialas, García-París, Ruiz & Recuero, 2020 | | | Spain: Andalucía:  Málaga: Arriate. | 43,961 | 437.1 | | 15,626 bp | |  | |
| SN1700F | Meloinae | | Lyttini | | *Oenas fusicornis* Abeille de Perrin, 1880 | | | Spain: Andalucía:  Sevilla: Gerena. | 83,642 | 780.4 | | 15,638 bp | |  | |
| APP15034B | Meloinae | | Lyttini | | *Lagorina sericea* (Waltl, 1835) | | | Spain: Andalucía:  Cádiz | 79,473 | 779.9 | | 15,631 bp | |  | |
| mel08233a | Meloinae | | Mylabrini | | *Hycleus dufourii* (Graells, 1849) | | | Spain: Extremadura:  Cáceres: Puerto de Honduras. | 15,895 | 242.9 | | 9,943 bp | |  | |
| SN1700Ca | Meloinae | | Mylabrini | | *Actenodia billbergi* (Gyllenhal, 1817) | | | Spain: Murcia: Puerto de Mazagón. | 91,026 | 868.5 | | 15,660 bp | |  | |
| SN1700G | Meloinae | | Mylabrini | | *Mylabris quadripunctata*  (Linnaeus, 1767) | | | Spain: Castilla-La Mancha: Toledo: 6 km West of Villacañas. | 51,972 | 477.3 | | 16,683 bp | |  | |
| APP15075 | Meloinae | | Mylabrini | | *Croscherichia paykulli* (Billberg, 1813) | | | Morocco: Souss-Massa: Chtouka Aït Baha: Biougra. | 81,058 | 854.3 | | 15,658 bp | |  | |
| mel08046 | Meloinae | | Eupomphini | | *Tegrodera latecincta* (Horn, 1891) | | | United States: California: Inyo County: Rudolph Road, 12 km Northeast Bishop, Highway 6. | 75,301 | 1016.4 | | 11,249 bp | |  | |
| mel06181 | Meloinae | | Eupomphini | | *Megetra punctata*  Selander 1965 | | | United States: Arizona: Cochise County: Portal. | 89,114 | 888.6 | | 15,640 bp | |  | |
| mel06011 | Meloinae | | Meloini | | *Meloe cavensis* Petagna, 1819 | | | Morocco: Casablanca-Settat: Benslimane:  Ouled Bahmad. | 30,884 | 416.6 | | 11,261 bp | |  | |
| mel81048 | Meloinae | | Meloini | | *Meloe mediterraneus* Müller, 1925 | | | Spain: Madrid: Colmenar de Oreja. | 89,759 | 932.1 | | 14,768 bp | |  | |
| mel08042a | Meloinae | | Meloini | | *Spastonyx nemognathoides* Selander, 1954 | | | United States: California: Mono County: Pumice Mill Road, 13.5 km Northeast of Bishop. | 176,026 | 1682.9 | | 15,627 bp | |  | |
| SN1700Ba | Meloinae | | Meloini | | *Physomeloe corallifer* (Germar, 1818) | | | Spain: Madrid: Manzanares. | 169,478 | 1793.4 | | 14,576 bp | |  | |
| mel06156b | Meloinae | | Epicautini | | *Epicauta tenella* (LeConte, 1858) | | | United States: California: San Bernardino County: Needles. | 201,771 | 1965.3 | | 15,708 bp | |  | |
| mel06211a | Meloinae | | Pyrotini | | *Pyrota palpalis* Champion, 1893 | | | United States: New Mexico: Valencia County: Los Lunas. | 29,627 | 368.8 | | 12,138 bp | |  | |
| melx11030 | Nemognathinae | | Horiini | | *Cissites auriculata*  (Champion, 1862) | | | Mexico: Jalisco: La Huerta: Chamela. | 94,578 | 969.9 | | 15,015 bp | |  | |
| mel06214 | Nemognathinae | | Nemognathini | | *Gnathium nitidum*  Horn, 1870 | | | United States: New Mexico: Valencia County: Los Lunas. | 254,647 | 2623.1 | | 14,991 bp | |  | |
| mel06252a | Nemognathinae | | Nemognathini | | *Zonitis atripennis*  Say, 1824 | | | United States: Utah: Iron County: Modena. | 243,961 | 3283.8 | | 11,240 bp | |  | |
| mel06167a | Nemognathinae | | Nemognathini | | *Zonitis (Parazonitis) tarasca borealis* Enns, 1956 | | | United States: Arizona: Cochise County: McNeal. | 207,785 | 2805.7 | | 11,238 bp | |  | |
| mel04038 | Nemognathinae | | Nemognathini | | *Leptopalpus rostratus* (Fabricius, 1792) | | | Spain: Andalucía: Sevilla: Alcalá de Guadaira. | 228,776 | 2234 | | 15,614 bp | |  | |
| mel08002 | Nemognathinae | | Nemognathini | | *Apalus guerini*  (Mulsant, 1858) | | | Spain: Madrid: Perales de Tajuña. | 101,300 | 976.5 | | 15,617 bp | |  | |
| **Genbank mt genomes of Meloidae** | | | | | | | | | | | | | | | |
| **Subfamily** | | **Tribe** | | **Species** | | **Locality** | **Reference** | | **Length** | | **GB accession number** | |  | | |
| Meloinae | | Epicautini | | *Epicauta chinensis* (Laporte, 1840) | | China: Shaanxi: Suide. | Du et al. (2016) | | 15,717 bp | | KP692789 | |  | | |
| Meloinae | | Epicautini | | *Epicauta gorhami* (Marseul, 1873) | | China: Shaanxi: Suide. | Du et al. (2017) | | 15,691 bp | | KX161854 | |  | | |
| Meloinae | | Epicautini | | *Epicauta tibialis* (Waterhouse, 1871) | | China: Guangdong: Beihai. | Du et al. (2017) | | 15,816 bp | | KX161855 | |  | | |
| Meloinae | | Epicautini | | *Epicauta aptera* (Kaszab, 1952) | | China: Nanchuan: Chongqing. | Jie et al. (2016) | | 15,645 bp | | NC031820 | |  | | |
| Meloinae | | Lyttini | | *Lytta caraganae* (Pallas, 1781) | | China: Guangdong: Beihai. | Du et al. (2017) | | 15,923 bp | | KX161859 | |  | | |
| Meloinae | | Mylabrini | | *Hycleus marcipoli* (Pan & Bologna, 2014) | | China: Guangdong: Beihai. | Du et al. (2017) | | 15,923 bp | | KX161857 | |  | | |
| Meloinae | | Mylabrini | | *Hycleus pharelatus* (Pallas, 1782) | | China: Guizhou: Luodian. | Du et al. (2017) | | 16,003 bp | | KX161858 | |  | | |
| Meloinae | | Mylabrini | | *Mylabris aulica* (Ménétriès, 1832) | | China: Inner Mongolia: Dongsheng. | Du et al. (2017) | | 15,758 bp | | KX161860 | |  | | |
| **Genbank mt genomes of Tenebrionoidea (outgroup)** | | | | | | | | | | | | | | | |
| **Family** | | **Species** | | **Country** | | **Reference** | **Length** | | **GB accession number** | |  | |  | | |
| Anthicidae | | *Formicomus*  La Ferté-Sénectère, 1849 | | South Africa | | Timmermans et al. (2015) | 10,593 bp | | JX412857 | |  | |  | | |
| Anthicidae | | *Omonadus floralis*  (Linnaeus, 1758) | | Czech Republic | | Timmermans et al. (2010) | 12,235 bp | | HQ232825 | |  | |  | | |
| Mordellidae | | *Mordella atrata* (Melsheimer, 1845) | | United States | | Cameron et al. (2009) | 15,540 bp | | FJ859904 | |  | |  | | |
| Mycetophagidae | | *Mycetophagus quadripustulatus*  (Linnaeus, 1761) | | Slovakia | | Timmermans et al. (2010) | 12,281 bp | | HQ232824 | |  | |  | | |
| Mycteridae | | *Stilpnonotus mexicanus* (Thomson, 1860) | | Belize | | Timmermans et al. (2015) | 11,447bp | | JX412811.1 | |  | |  | | |
| Oedemeridae | | *Ischnomera cyanea* (Fabricius, 1792) | | Czech Republic | | Timmermans et al. (2015) | 12,524 bp | | JX412790 | |  | |  | | |
| Oedemeridae | | *Oedemera virescens*  (Linnaeus, 1767) | | Czech Republic | | Timmermans et al. (2010) | 10,653 bp | | HQ232826.1 | |  | |  | | |
| Tenebrionidae | | *Asbolus verrucosus* (LeConte, 1851) | | ------ | | Rider (2016) | 15,828 bp | | NC027256.1 | |  | |  | | |
| Tenebrionidae | | *Tenebrio molitor* (Linnaeus, 1758) | | China? | | Li-Na & Cheng-Ye (2014) | 15,785 bp | | KF418153 | |  | |  | | |
| Tenebrionidae | | *Tenebrio obscurus* (Fabricius, 1792) | | China? | | Bai et al. (2018) | 15,771 bp | | NC037196.1 | |  | |  | | |
